# Integrative analysis of genomic and transcriptomic data informs precancer progression in the pancreas

**DOI:** 10.1101/2025.11.03.686234

**Authors:** Kathleen Noller, Jiaying Lai, Daniel Lesperance, Ricky S. Adkins, Ahmed Elhossiny, Paola A. Guerrero, Kimal I. Rajapakshe, Anirban Maitra, Michelle Giglio, Anup Mahurkar, Owen White, Marina Pasca Di Magliano, Michael F. Ochs, Luciane T. Kagohara, Laura D. Wood, Rachel Karchin, Elana J. Fertig

## Abstract

Pancreatic ductal adenocarcinoma (PDAC) arises from heterogeneous precursor lesions, including intraductal papillary mucinous neoplasms (IPMNs), but the features distinguishing indolent from progressive lesions remain unclear. We performed an integrative analysis of transcriptomic, genomic, and microenvironmental profiles of IPMNs to define multi-omic phenotypes. Using transfer learning, we projected IPMN-derived transcriptional programs onto spatial transcriptomic datasets from IPMNs and pancreatic intraepithelial neoplasias (PanINs). We identified two major phenotypes: one associated with cancer-associated fibroblasts and epithelial-to-mesenchymal transition, shared across IPMN, PanIN, and PDAC; and a second, glycolysis-enriched phenotype with a unique somatic mutation profile specific to IPMN. Spatial mapping further revealed grade-specific enrichment of transcriptional programs and distinct interactions with stromal and immune subtypes, underscoring the role of the precancer microenvironment in progression. These findings establish multi-omic phenotypes that unify genetic, transcriptional, and microenvironmental heterogeneity, providing a framework for distinguishing progressive from indolent precancers and a web-based public atlas for future exploration of these data and transcriptional phenotypes.

## Introduction

Pancreatic ductal adenocarcinoma (PDAC) is a deadly disease with an overall five-year survival rate of approximately 10% despite recent improvements in survival in a subset of patients upon surgical resection of the tumor^1^. Pancreatic cancer has the lowest survival rate of any major organ cancer and is the third leading cause of cancer mortality in the United States^2^. PDAC evolves from precancerous lesions, most commonly pancreatic intraepithelial neoplasia (PanIN) and intraductal papillary mucinous neoplasm (IPMN). Indolent and low-grade precancerous lesions are prevalent in the healthy patients regardless of age, with only a subset progressing to PDAC^3, 4^.

PDAC patients suffer chemotherapy resistance and late diagnoses with a high incidence of metastases^5^, however the early detection of IPMN lesions is possible on abdominal imaging, presenting an opportunity for early intervention^6^. IPMNs are highly heterogeneous: each lesion contains multiple subclones with distinct molecular alterations, which undergo selection preceding the development of select lesions into high-grade IPMN and PDAC^7^. The impact of neoplastic cell phenotypes and context of the microenvironment in this selection process remains unknown.

We also hypothesize that there is a relationship between transcriptional patterns and both the genomic profile of the lesions and the precancerous microenvironment (PME) that are associated with progression. In advanced PDAC, the cellular phenotypes associated with progression have been well studied. For example, two well-known transcriptional subtypes characteristic of the PDAC epithelium correlate with patient survival and progression to metastasis, including notably the “classical” and “basal-like” subtypes from Moffitt et al.^8^. Two other PDAC-associated epithelial transcriptional signatures have also been defined by Collisson et al.^9^, which delineates classical, exocrine, and quasi-mesenchymal subtypes, and by Bailey et al.^10^, which delineates squamous, progenitor, immunogenic, and ADEX (aberrantly differentiated endocrine exocrine) subtypes correlated with histopathological features. Moffitt et al. have also performed transcriptional subtyping of the stromal and immune compartments in PDAC^8^. Studies of primary and metastatic PDAC have already demonstrated the critical role of the fibrotic, metabolic, and immunosuppressive microenvironments in PDAC invasion^11^ and regulating transitions between transcriptional phenotypes in tumor cells^12,13^. However, application of these subtypes to pancreatic precancer has demonstrated that precancer lesions are predominantly classical, however transcriptional variation and genomic subclonal heterogeneity in these precancerous lesions have been observed^14,15^. There is a need to develop new pancreatic precancer-specific subtypes with the goal of distinguishing indolent and progressive lesions. However similar analyses in IPMNs or with precancer-specific patterns has not yet been performed.

In this study, we define multi-omic subtypes of IPMN lesions through unsupervised pattern detection of matched genomic and bulk transcriptomic data from low and high-grade IPMN lesions^16^. We perform transfer learning^17^ of phenotype-associated transcriptional patterns onto IPMN^18^ and PanIN^14^ spatial transcriptomic data to infer PME traits and histological features associated with each phenotype. We identify two IPMN-enriched phenotypes with distinct somatic mutation profiles, one of which is IPMN-specific while the other is similarly enriched in PanINs, where it correlates with other cancer-associated fibroblast (CAF)-associated transcriptional features. These phenotypes provide a deeper characterization of pancreatic precancer transcriptional and genomic heterogeneity and the relationship between PDAC, IPMN, and PanIN subtypes. Beyond these subtypes, we collate these data into a comprehensive pancreatic precancer atlas in a new extension of the user-friendly web-based portal gEAR^19^, called Cancer-gEAR, to enable broad characterization of precancerous lesions for custom analyses and querying our inferred transcriptional signatures in additional datasets.

## Methods

### Bulk RNA-seq analysis

We obtained whole transcriptome Smart-3Seq data from *n =* 24 patients with *n =* 252 samples with histological areas annotated as either normal ductal (ND), normal acinar (Ac), low or high grade IPMN, or PDAC from Semaan et al.^16^. Raw HTSeq counts files were obtained for each sample and samples with 3^rd^ quantile count values of zero (*n =* 43 samples) were removed from further analysis. Size factor estimation, dispersion estimation, and variance stabilizing transformation (VST) were performed in DESeq2 (version 1.44.0)^20^.

### Classification of transcriptional data

A rank-based classifier (https://github.com/rmoffitt/pdacR)^21^ was applied to the VST-transformed counts matrix to classify RNA-seq samples according to basal and classical PDAC transcriptional subtypes. While a separate tool (CMScaller^22^) was used in Semaan et al. to classify samples based on expression profiles of marker genes, we elected to apply the classifier developed in the original Moffitt study^8, 21^ to compute the per-sample probability of basal-like class membership. We also obtain marker gene lists from two other published PDAC-associated epithelial signatures: Collisson et al.^9^ and Bailey et al.^10^. Samples were annotated based on their maximum gene set expression of a given marker gene list.

### Whole Exome Sequencing data analysis

Whole-exome sequencing (WES) data (accessed via dbGaP accession number: phs002225.v3.p1) were pre-processed using Trim Galore (version 0.6.10) for quality and adapter trimming, then aligned to the human reference genome (hg38) using bwa mem (version 0.7.17). The GATK (version 4.4.0.0)^23^ best practices pipeline was applied for data pre-processing and somatic short mutation discovery with Mutect2^24^. Mutations were retained only if, in at least one sample, total reads ≥ 50, alternate reads ≥ 20, and VAF ≥ 0.10. Any mutations labeled as “weak evidence” by GATK’s Mutation Quality Score Recalibration (VQSR) were removed. Visual inspection was performed with IGV (version 2.16.0)^25^. Mutations were annotated with OpenCRAVAT (version 2.4.1)^26^. Copy number alterations (CNAs) were called using FACETS (version 0.6.2)^27^.

### Subclonal analysis

Genomics-based subclonal analysis was performed with PICTograph (verson 1.3.2)^28^ with default settings and MCMC parameters *n.iter*=50,000, *n.burn*=10,000. The sample presence feature was not applied. For each patient, only one sample per lesion type was included when multiple samples were available. Previously reported IPMN/PDAC driver mutations were highlighted on the inferred tumor evolutionary trees^29, 30^

### Pattern detection analysis

CoGAPS, a non-negative matrix factorization-based tool, was used for latent space learning on both an individual patient basis and a population basis (version 3.24.0)^31^. CoGAPS was run with the following parameters: sparse optimization enabled, 10,000 iterations, output frequency of 1000, sparsity parameter alpha of 0.01, and 9 patterns. The optimal number of patterns for this dataset was chosen from an elbow plot of mean chi squared statistic versus number of patterns and from correlative analysis of pattern matrix values for different CoGAPS runs on the same dataset. CoGAPS was run on the VST-transformed counts matrix of RNAseq data. To obtain rankings of features for a given pattern, the function *patternMarkers* was used. To run GSEA on a given pattern, the Molecular Signatures Database was used to obtain Hallmark reference pathways (msigdbr, version 7.5.1)^32^ and the CoGAPS function *getPatternGeneSet* was used to find enriched gene sets with the enrichment method.

### Integrative analysis

Correlation-based clustering was performed on the pattern weights matrix using the *agnes* function from the cluster package (version 2.1.8.1) in R. Pattern 8, which was determined to be associated with read count, was excluded from clustering. Five clusters were obtained containing a minimum number of 3 individual set contributions. Fisher’s exact test with Bonferroni adjustment of p-values was used to compare mutation distribution across clusters, excluding the normal duct-specific cluster which contains no somatic mutations. Somatic mutations for which integrative analysis was performed were chosen based on occurrence in previous studies of IPMN/PDAC driver genes^29, 30^. OpenCRAVAT was used to obtain CHASMplus scores of somatic mutations^26^. CHASMplus q-values were computed from Benjamini-Hochberg analysis, since pre-computed CHASMplus q-values are not available on OpenCRAVAT. The COSMIC database^33^ was used to obtain the number of occurrences of each mutation across multiple studies.

### Deconvolution analysis

Deconvolution of bulk RNA-seq data was performed in InstaPrism (version 0.1.6)^34^. An annotated scRNA-seq dataset containing tissue from the adult healthy human pancreas was used as reference^4^. Estimated cell type fractions were obtained per query sample.

### Spatial transcriptomic data analysis

Analysis of publicly available Visium 10X genomics data containing IPMN samples from Sans et al.^18^ or PanIN samples from Bell et al.^14^ was performed in Seurat (version 5.1.0)^35^. ProjectR (version 1.20.0)^17, 36^ was used to project feature weights of transcriptional patterns onto the spatial transcriptomic query dataset. To identify putative areas of interaction of two distinct cell types, we used SpaceMarkers (version 1.0.0)^37^ with author-provided RCTD-derived cell type scores^18^ or author-provided CODA-derived cell type scores^14^ as input to identify hotspots and areas of interaction in differential expression (DE) mode. Patterns were skipped if no significant interactions with any cell type were identified.

### Statistics

Pearson correlations were obtained using the *cor.test* function in R. Differences in pattern weight across multiple groups were computed using ANOVA with post-hoc Tukey’s HSD or with the Kruskal-Wallis test if assumptions of ANOVA were violated (*aov*, *TukeyHSD*, and *kruskal.test* in the base R stats package, version 4.4.0).

### Cancer gEAR platform

The Pancreas Cancer gEAR (cancer.umgear.org) instance was cloned in Google Cloud from the Gene Expression Analysis Resource^19^. Spatial transcriptomic data were loaded into the Pancreas Cancer gEAR and fed into the projection analysis tool. The projection analysis in the Pancreas Cancer gEAR instance utilizes projectR (verision 1.23.2)^17^ on multiple serverless instances for processing. Each dataset is broken into smaller chunks, and for each chunk, the pattern matrix and relevant arguments are passed to ProjectR. Each result is then sent back to the Pancreas Cancer gEAR server where the results are concatenated to create a dataset-level result.

## Results

### Population-wide, mutation-level genomic heterogeneity exceeds per-patient, subclone-level heterogeneity

The goal of our study is to associate genomic variation with transcriptional phenotypes that regulate dysplastic progression from a range of pancreatic precancer datasets (Table 1). We begin our analysis with a cohort of matched transcriptomic (bulk RNA-sequencing) and genomic (whole exome sequencing) data from low and high grade IPMN, PDAC, and normal ductal tissue^16^. Unlike the other precancer datasets examined, this dataset does not offer spatial information yet provides matched genomic and transcriptomic data for the inference of multi-omic phenotypes. We began with a mutational analysis of this data to assess whether we could obtain cohort-wide genomic profiles that could then be related to transcriptional patterns in the data. We performed subclonal inference and somatic mutation calling to identify total of 488 unique coding mutations across all patients. As expected, we observed little patient-to-patient overlap of somatic mutations, with only 7 mutations, six of which are located in known PDAC driver genes or genes with known roles in PDAC progression (*KRAS*, *GNAS*, *KLF4,* and *MYCN*), shared across 2 or more patients **(Figure 1A)**. Among these mutations are some of the most common mutational drivers of PDAC progression such as *KRAS* G12D (p.Gly12Asp), *KRAS* G12V (p.Gly12Val), and *KRAS* G12R (p.Gly12Arg). In PDAC, the most predominant *KRAS* mutation sites occur at G12D (45%), G12V (35%) and G12R (17%)^38^.

**Figure 1.**
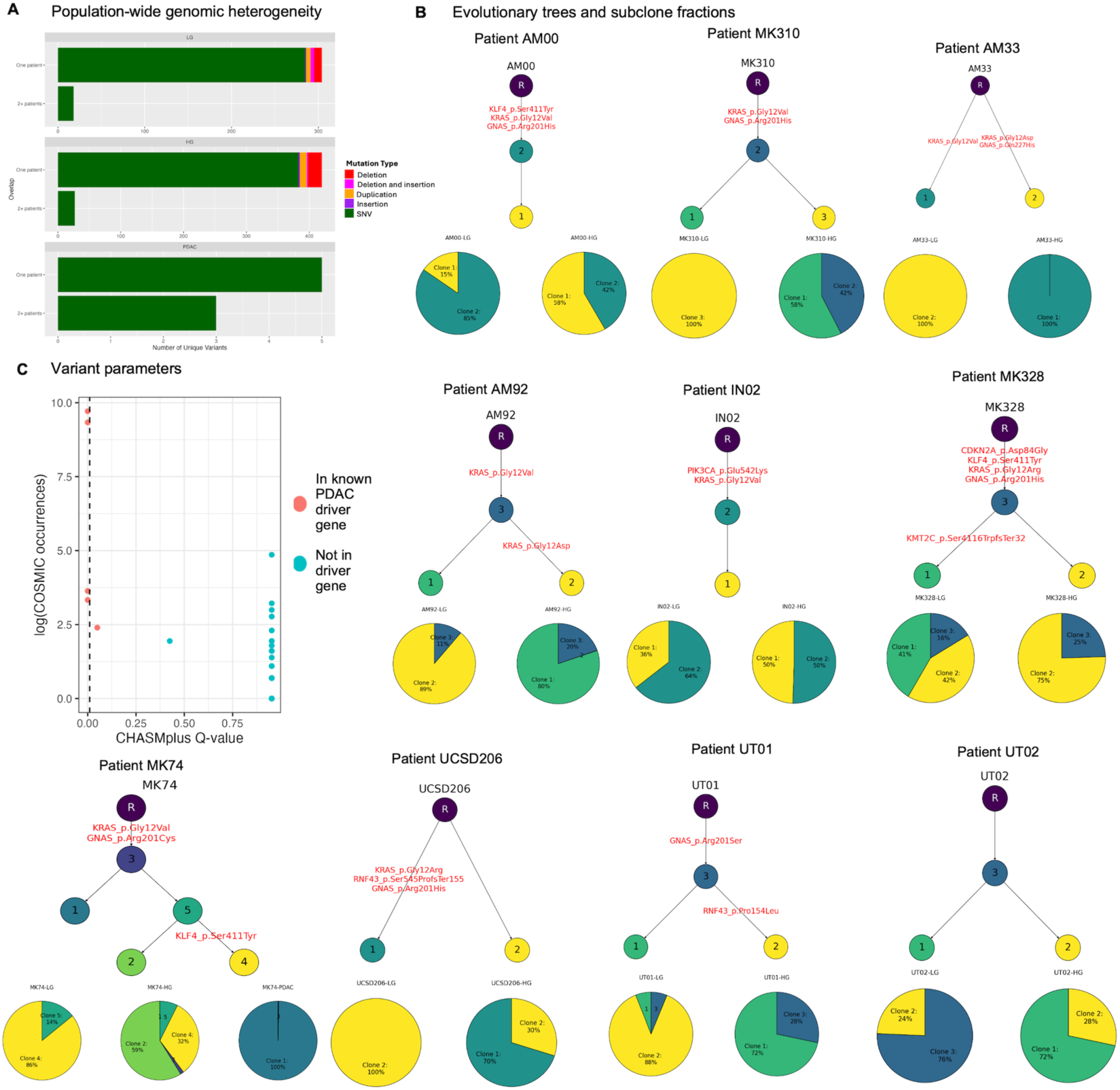
Comparison of genomic within-patient and population-wide heterogeneity

To perform per-patient genomics analysis, we use the Bayesian hierarchical model PICTograph to identify patient-specific subclones and to reconstruct probable ancestral relationships between subclones^28^. This tool estimates per-patient genomic heterogeneity, as measured by number of subclones, or genetically distinct populations within a tumor, per patient, as it infers clonal evolution. We depict ancestral relationships between subclones as well as subclone proportion per tissue type (low or high grade IPMN, and PDAC) for every patient with greater than one WES sample and with greater than five mutations **(Figure 1B).** Within-patient genomic heterogeneity, represented by the number of distinct genomic subclones per patient, ranges from 2 to 5 genomically distinct subclones per patient. Additionally, evolutionary tree geometry differs by patient, with both simple branched and linear trajectories observed, meaning that multiple genetically distinct subclones may arise from a single subclone of origin and suggesting that their evolution may be driven by distinct genomic profiles and possibly by distinct microenvironmental factors. These results demonstrate that per-patient genomic heterogeneity, while present at both the individual mutation level and the subclone level, is much lower than population-wide genomic heterogeneity, and that drivers of dysplastic progression generalizable to a population may be limited to common and well-known oncogenic mutations in addition to microenvironmental or transcriptomic factors.

We hypothesize that only a small fraction of the mutations that distinguish these precancer subclones are drivers. To test this hypothesis, we obtain quantitative measures for individual mutations from the CHASMplus^39^ PAAD and COSMIC databases^33^: CHASMplus q values (where q < 0.01 is suggestive of a driver) and number of occurrences of a mutation in the COSMIC cancer-associated mutation database. We pay particular attention to mutations located in the coding region of known PDAC driver genes: *ARID1A, ARID2, CDKN2A, GNAS, KMT2C, SMAD4, SMARCA4, RNF43, TGFBR2, TP53, KRAS, SF3B1, KLF4, PIK3CA*, and *BRAF*^29, 30^.

Out of 488 mutations, a total of 54 mutations across all samples, 8 of which are located in known PDAC driver genes, contain entries in both the CHASMplus and COSMIC databases. Amongst these, all except two (KLF4 c.1232C>A and GNAS c.681G>T) of the mutations located in known PDAC driver genes contained both a low CHASMplus q value and a relatively high number of COSMIC occurrences **(Figure 1C)**. These metrics suggest that the true driver mutations of PDAC are contained within our list of mutations in known driver genes.

### Precancerous transcriptional subtypes define cohort-wide phenotypes

We next sought to quantify transcriptional states in IPMNs from the corresponding RNA-seq data in the Semaan et al. cohort^16^. While Semaan et al. performed subtype classification according to previously defined PDAC epithelial transcriptional subtypes, we developed a comprehensive pipeline to infer IPMN-specific subtypes in our own study so that we could apply it to all precancer datasets we sought to study. Before doing so, we define IPMN transcriptional subtypes against PDAC-specific subtypes already present in the literature. We classify the RNA-seq samples from Semaan et al. according to well-established epithelial PDAC transcriptional subtypes associated with significant differences in patient survival—basal-like and classical—using the rank-based classifier from the original manuscript^8^. Consistent with Semaan et al., analysis on this cohort demonstrates a high proportion of “classical” samples and a low proportion of “basal-like” samples according to both Moffitt and Collisson definitions (**Supplemental Table 1**). Methodological differences are responsible for most differences between our results and those of Semaan et al; for example, most samples (74%) return “no matched classification” for the Moffitt categories in the Semaan et al. study whereas the classifier we apply returns basal-like probability, not a per-sample labeling. Our analysis reveals that both low and high grade IPMN samples universally exhibit very low basal-like probability, with a maximum of 0.056 out of a possible 1.0 **(Figure 2A)**. In both our study and Semaan et al, there is little variation in transcriptional subtypes between low grade and high grade IPMN lesions: for the Collisson gene sets, all but two IPMN samples classify as Collisson classical subtype and none classify as the quasi-mesenchymal subtype **(Figure 2A).** For the Bailey gene sets, IPMN samples classify predominantly as a mixture of immunogenic and progenitor subtypes, the former likely from contamination of immune cells, however those samples do not separate cleanly in reduced dimensional space **(Figure 2A).** The cluster of normal ductal samples marked as Bailey ADEX or Collison exocrine subtypes fit with the prevailing hypothesis that ADEX and exocrine-like subtypes are the results of contamination from normal acinar tissue^40^. These results suggest that the Collisson and Bailey subtypes may, like the Moffitt subtypes, hold less utility for IPMN subtyping than for PDAC, suggesting that *de novo* inference of cell state transitions in this IPMN cohort is needed.

**Figure 2.**
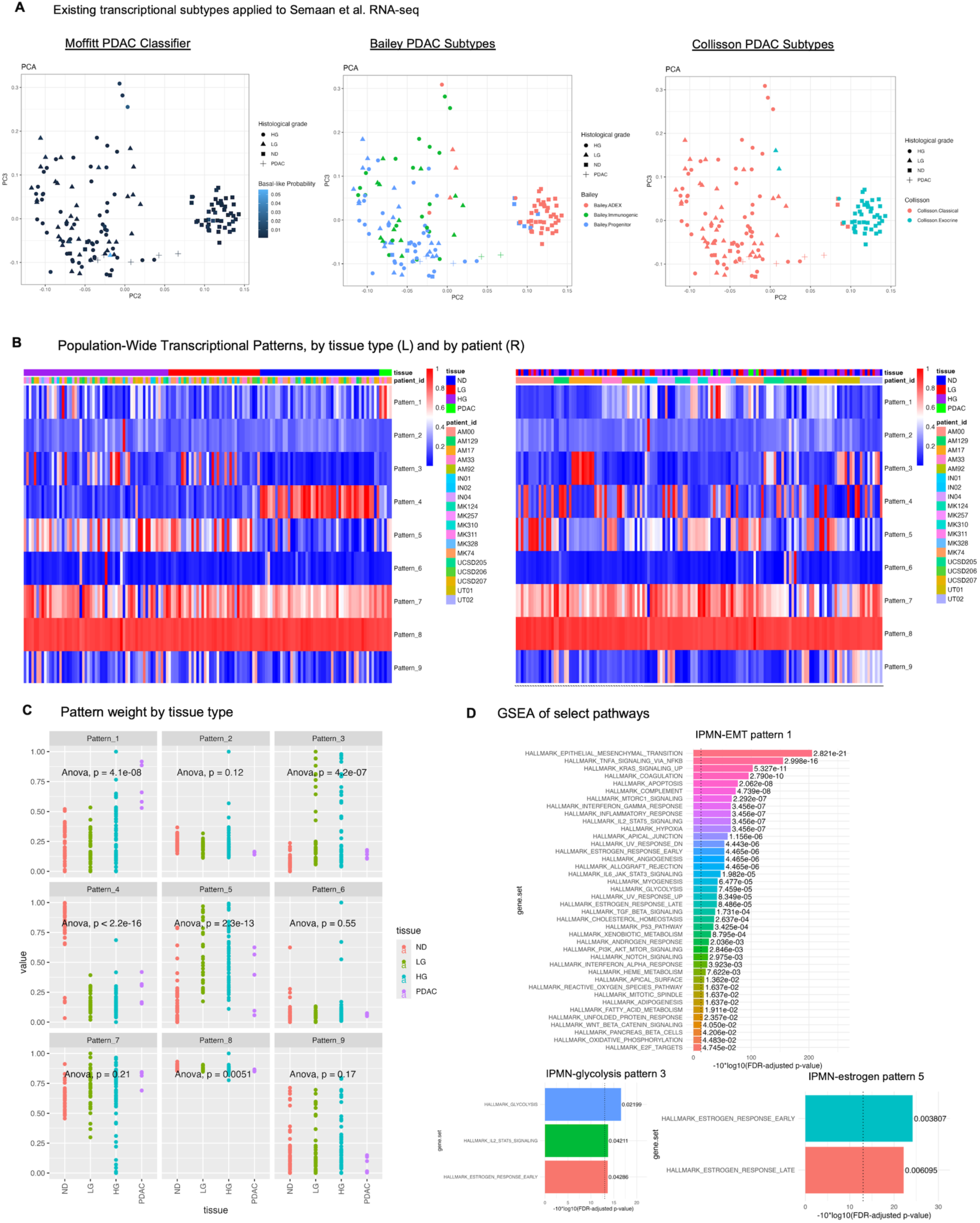
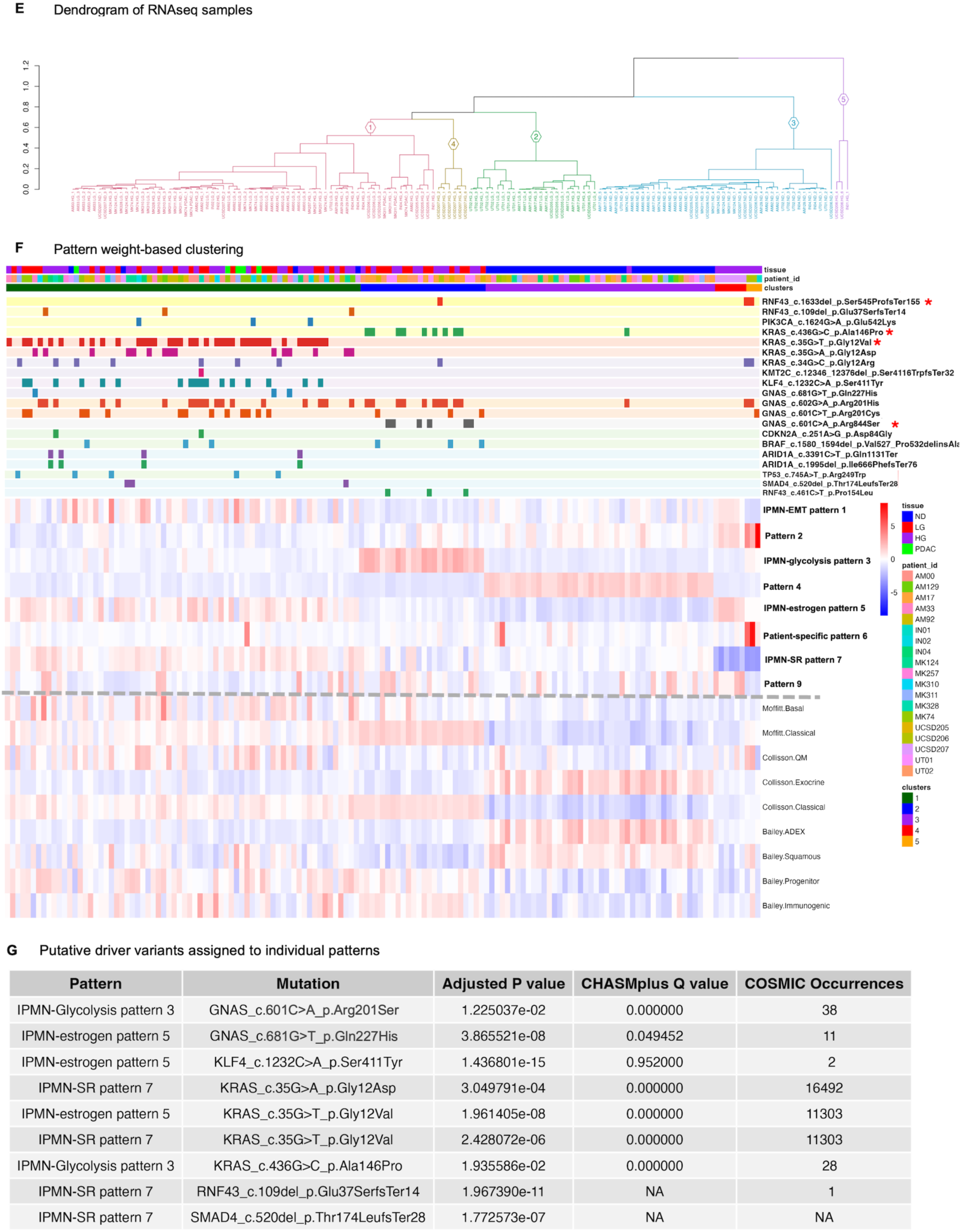
Population-wide transcriptional patterns transcend per-patient genomic heterogeneity

To investigate whether the consistency of transcriptional subtypes between IPMN grades is due to the classification or whether additional definition of transcriptional phenotypes learned in IPMN could distinguish grade, we perform pattern detection analysis on the transcriptional data. We apply CoGAPS, a Bayesian non-negative matrix factorization tool^31, 41^, to identify unique transcriptional patterns present in the data. We ran CoGAPS on RNAseq samples from all patients containing matched transcriptional and genomic data and on samples derived from all ductal epithelial tissue types (normal duct, low and high grade IPMN, and PDAC).

Optimization of number of patterns **(Supplementary Figure 1A-B)** found 9 patterns in the RNA-seq data, one of which was associated with baseline expression and read depth, leaving 8 patterns for downstream analysis **(Figure 2B)**.

Of these 8 patterns, two patterns are sample-specific and patient-specific, respectively: pattern 2 is highly weighted in one sample, and pattern 6 is highly weighted in one patient as compared to others (patient UCSD206), however all other transcriptional phenotypes are found across multiple patients. If we represent population-wide transcriptional heterogeneity as the number of patterns needed to represent the transcriptional complexity of the dataset, it is lower than the number of shared genomic mutations across more than one patient and offers an opportunity to identify patterns unifying samples from different patients. We identify four tissue-specific patterns, with pattern 4 clearly enriched in normal duct as compared to all other tissue types (p < 2.2e-16, ANOVA), pattern 1 enriched in PDAC samples as compared with all other tissue types (*p* = 4.1e-08), and patterns 3 and 5 enriched in LG and HG IPMN as compared with normal ductal epithelium (*p* = 4.2e-7 and *p =* 2.3e-13, ANOVA with Tukey post-hoc HSD) **(Figure 2C).** Of note, there is no transcriptional pattern which separates out low grade from high grade IPMN, which corroborates the lack of separation of LG and HG IPMN samples in reduced dimensional space **(Figure 2A).**

We therefore turn our attention to patterns 1, 3 and 5 as enriched in PDAC and/or IPMN. Gene set enrichment analysis of these patterns reveals expected upregulation of pathways associated with PDAC in pattern 1, including epithelial-to-mesenchymal transition, TNF-alpha signaling via NF-kB, and KRAS upregulation (*p adjusted* = 2.821e-21, 2.998e-16, and 5.327e-11, respectively). Pattern 3 is enriched in glycolysis, IL2-STAT5 signaling, and early estrogen response (*p adjusted* = 0.2199, 0.04211, and 0.04286, respectively) and pattern 5 is enriched in estrogen response pathway activity (*p adjusted = 0.003807* and 0.006095 for estrogen response early and late Hallmark pathways) **(Figure 2D).** We will refer to these transcriptional patterns for clarity as “IPMN-EMT pattern 1,” “IPMN-glycolysis pattern 3,” and “IPMN-estrogen pattern 5” throughout the remainder of this study. Gene set enrichment analysis results for other patterns whose weights are not significantly different by tissue type are provided in **Supplementary Figure 2A-D.** Pattern 7, for example, while not found to be significantly enriched in IPMN or PDAC samples, exhibited GSEA terms associated with cell survival and stress response such as TNF-alpha signaling, hypoxia, p53, and MTORc1 signaling and may represent a pan-epithelial stress response program (henceforth referred to as “IPMN-SR pattern 7”) (**Supplementary Figure 2C).**

To partition samples into categories based on expression of identified transcriptional programs, we perform correlation-based clustering of RNA-seq samples based on pattern weights of the 8 CoGAPS patterns **(Figure 2E)**. As expected, the normal ductal samples localize to a single cluster (cluster 3, deemed the “normal duct (ND) cluster”) in which samples are enriched in the phenotype represented by transcriptional pattern 4. Interestingly, IPMN samples do not form subclusters based on histological grade, with both low and high grade IPMN dispersed throughout clusters 1 and 2. While all samples contain low basal-like probability per the Moffitt classifier and are therefore classified as belonging to the PDAC “classical” subtype, we do observe significant differences in classical gene set score between clusters, though not in basal-like gene set score, upon comparison of IPMN and PDAC-enriched clusters. Classical gene set score is significantly higher in cluster 2 than clusters 1, 4, or 5 (*p adjusted < 0.05, Kruskal-Wallis with Dunn post-hoc,* **Figure 2F),** suggesting that while all samples are classified as “classical,” there still exist differences in the magnitude of gene expression of classical marker genes.

### Pattern-based IPMN clusters contain unique somatic mutation profiles of known PDAC drivers

While the Semaan et al. cohort contains matched transcriptomic and genomic data, RNAseq and WES samples from the same patient and tissue type were taken from adjacent regions via laser capture microdissection per the original authors, with multiple RNA-seq replicates per tissue type often taken yet only one WES sample per tissue type. It is not guaranteed that the RNA-seq and WES samples contain identical somatic mutation profiles, thus even while we assume the same mutation profile for samples from the same patient and tissue type, our analysis is focused at the sample level rather than the subclone level.

Following our clustering analysis above, we find that IPMN samples segregate to clusters 1, 2, 4, and 5, each of which is represented by a unique pattern or combination of transcriptional patterns. For example, IPMN-glycolysis pattern 3 is highly specific for cluster 2, while samples with high weights of patterns IPMN-EMT pattern 1, IPMN-estrogen pattern 5, and IPMN-SR pattern 7 localize predominantly to cluster 1. PDAC samples are located exclusively in cluster 1. We therefore name these clusters as the IPMN-PDAC cluster (cluster 1) and the IPMN-P3 cluster (cluster 2). There is one patient-specific cluster (cluster 4), which contains only high grade IPMN (n = 9 samples), and there is one small cluster (n = 3) containing HG IPMN from two different patients which is associated with high weights in patterns 2 and 6 **(Figure 2F)**. We hypothesize that the IPMN-PDAC and IPMN-P3 clusters represent two generalizable transcriptional subtypes of IPMN, or perhaps of pancreatic precancer more broadly, which are associated with different genomic profiles but not with histological grade or classical or basal-like transcriptional phenotype.

To determine whether transcriptional patterns are associated with the distribution of putative genomic drivers of PDAC, we focus our analysis on the mutations contained in known PDAC driver genes. There are a total of 20 mutations located in these genes; six are deletions and the rest are single nucleotide mutations (SNVs). Of note, because of the high population-wide genomic heterogeneity and little patient-to-patient overlap of specific mutations, only six of these mutations are present in two or more patients (the other mutations are assumed to be present across replicates from the same patient and tissue type): *GNAS* c.601C>T (patients MK74, UCSD205, and IN01 in the IPMN-PDAC and IPMN-P3 clusters and in cluster 5), *GNAS* c.602G>A (patients MK310, AM129, AM17, MK328, AM00, and UCSD206 in the IPMN-PDAC and IPMN-P3 clusters and in cluster 5), *KLF4* c.1232C>A (patients MK74, MK328 and AM00 in the IPMN-PDAC cluster), *KRAS* c.34G>C (patients MK311, MK328, and UCSD206 in the IPMN-PDAC and IPMN-P3 clusters and in cluster 5), *KRAS* c.35G>A (patients AM33, AM129, AM92, and MK124 in the IPMN-PDAC cluster only), and *KRAS* c.35G>T (patients AM33, AM92, MK74, MK310, IN02, IN04, and AM00 in the IPMN-PDAC cluster only). While samples in the IPMN-PDAC cluster contain mutations that are both unique to that cluster and present across multiple patients, all mutations unique to the IPMN-P3 cluster are also unique to a single patient, therefore the generalizability of its genomic profile is limited. Upon analysis of mutation distribution amongst our IPMN clusters, we find that one *GNAS* SNV (c.601C>A), two *KRAS* SNVs (c.35G>T and c.436G>C), and one *RNF43* deletion (c.1633del) each have a significantly different distribution amongst clusters (*p adjusted* < 0.05, Fisher’s exact test with Bonferroni adjustment). None of these mutations are significantly differently distributed if samples are grouped by tissue type (LG IPMN, HG IPMN, or PDAC) as opposed to cluster assignment. If we bin individual mutations by gene name, then we report a significant difference only in the occurrence of *GNAS* and *KRAS* mutations by cluster (excluding the ND cluster, *p adjusted* < 0.05 for both comparisons, **Supplementary Figure 2E**). These results indicate that there exists some significant relationship between the occurrence of select mutations in known PDAC driver genes and the transcriptional phenotypes defined by our pattern weight-based clustering which is independent of tissue type and IPMN grade.

To assess whether individual transcriptional patterns themselves were associated with individual mutations, we binned RNA-seq samples by whether or not they contained a given genomic mutation and compared the weight of a given pattern in samples containing the mutations versus not. Mutations were assigned to one or more patterns if samples with versus without the mutation exhibited significantly higher pattern weight.

Four transcriptional patterns, three of which are enriched in dysplastic or cancerous tissue, were assigned a total of 202 mutations: IPMN-EMT pattern 1, IPMN-glycolysis pattern 3, IPMN-estrogen pattern 5, and IPMN-SR pattern 7. Among all of these mutations, we still observe that only the mutations in known PDAC driver genes contain low CHASMplus q values indicative of driver status or high COSMIC occurrences. Nine of these mutations were assigned to one of the four transcriptional patterns above **(Table 2G, Supplementary Table 2)**. These results support the idea that transcriptional phenotypes are associated with many mutations, only a handful of which are known to be important for progression to PDAC.

Given that mutations distinguished only a subset of the clusters of transcriptional patterns, we next sought to determine if any were mediated by the microenvironment. Deconvolution of the bulk RNA-seq data using a scRNA-seq reference dataset from healthy adult human pancreas tissue^4^ in InstaPrism was performed to assess how cell type fraction differences of samples in the original cohort might contribute to the differences observed in transcriptional programs. First, we perform a cluster-based analysis and find that the estimated fibroblast fraction in cluster 1 is the lowest amongst all clusters, despite the enrichment of IPMN-EMT pattern 1 in cluster 1, suggesting that this IPMN-EMT pattern underlying cluster 1 derives from the epithelial cells themselves, not from contaminating fibroblasts in the bulk sample **(Supplementary Figure 2F).** We then analyze differences in deconvolved fibroblast fraction with transcriptional pattern weight. Upon correlative analysis of estimated fibroblast fraction with transcriptional pattern weight in each tissue type, many patterns show a negative or non-significant Pearson correlation to fibroblast fraction across all tissue types (IPMN-glycolysis pattern 3, pattern 4, IPMN-estrogen pattern 5, patient-specific pattern 6, IPMN-SR pattern 7, and pattern 9). However, IPMN-EMT pattern 1 is the only pattern to exhibit a statistically significant, positive correlation with estimated fibroblast fraction in high grade IPMN (R = 0.66, p = 3.0e-8) but not in normal epithelium, low grade IPMN, or PDAC **(Supplementary Figure 2G).** IPMN-EMT pattern 1 is only one of three transcriptional patterns which are each highly weighted in cluster 1, thus we hypothesize that the EMT signal contributing to this phenotype from IPMN-EMT pattern 1 specifically may arise from physical proximity of IPMN epithelium to fibroblasts.

### Transfer learning analysis in spatial transcriptomic data reveals IPMN grade-specific transcriptional programs

A limitation of the previous dataset is that while the RNAseq and WES samples from the same patient and tissue type were taken from adjacent regions via laser capture microdissection per the original authors, it is not guaranteed that these samples contain identical somatic mutation profiles. To interrogate microenvironmental contributors to our transcriptional patterns and to interrogate the spatial landscape of these patterns, we perform transfer learning of the feature weights of each pattern to a spatial IPMN dataset. The results of CoGAPS run on the RNAseq samples was projected using ProjectR^17, 36^ onto the epithelial spots of a spatial transcriptomic dataset (Visium 10X) containing low grade IPMN, high grade IPMN, and PDAC lesions^18^. Spots were annotated by the original authors using Robust Cell Type Deconvolution (RCTD)^42^ and by spatial location relative to the neoplastic epithelium (epilesional, or overlapping with neoplastic epithelium, juxtalesional, or surrounding the neoplastic epithelium, or perilesional, distal to the juxtalesional region) **(Figure 3A)**.

**Figure 3.**
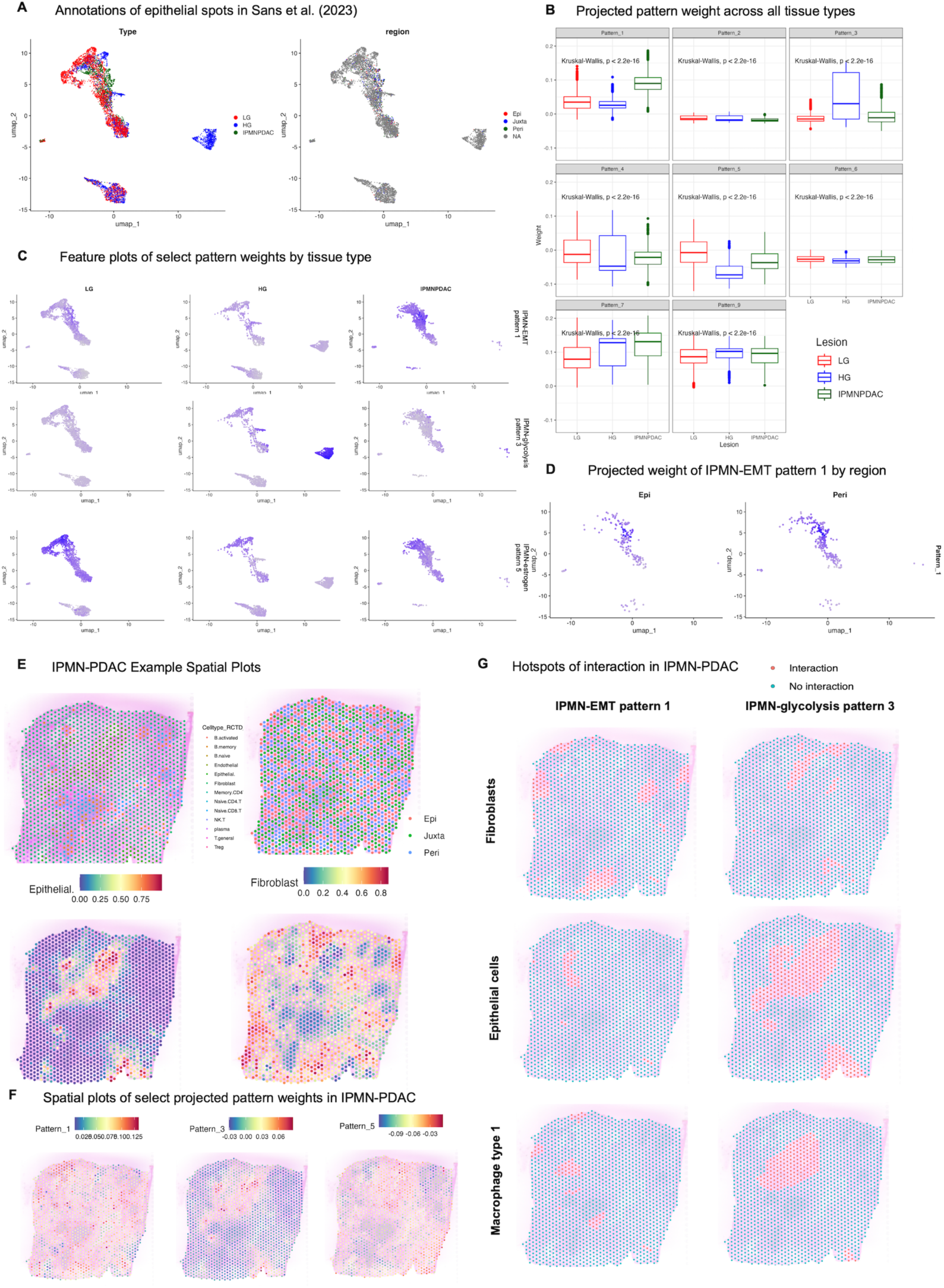
Transcripttional patterns are associated with microenvironmental features in IPMN

To determine how transcriptional patterns relate to microenvironment, we first examine how projected pattern weights differ by tissue type in the spatial data (LG IPMN, HG IPMN, or IPMN-PDAC) **(Figure 3B)**. IPMN-EMT pattern 1, IPMN-glycolysis pattern 3, and IPMN-estrogen pattern 5, which each exhibited differing weights by tissue type in our original cohort, also show significantly different projected weights in this spatial query dataset **(Figure 3C)**. IPMN-EMT pattern 1 has a significantly higher weight in IPMN-PDAC than in either LG or HG IPMN (p < 2.2e-16, Kruskal-Wallis, p adjusted = 0 for HG vs. IPMN-PDAC, p adjusted = 0 for LG vs. IPMN-PDAC, Dunn post-hoc test with Bonferroni adjustment); this confirms the elevated weight observed for this pattern in PDAC samples from our original cohort. IPMN-glycolysis pattern 3 is most highly weighted in HG IPMN as compared to LG or IPMN-PDAC (p < 2.2e-16, Kruskal-Wallis, p adjusted = 1.41e-103 for HG vs. IPMN-PDAC, p adjusted = 1.07e-276 for HG vs. LG, Dunn post-hoc test with Bonferroni adjustment). In our original cohort, none of the patterns delineated low from high grade IPMN; IPMN-glycolysis pattern 3 was highly specific to the IPMN-P3 cluster but was enriched in both high and low grade IPMN. Perhaps viewing this pattern in the context of a spatial dataset with increased resolution has revealed its importance in high-grade IPMN specifically. Finally, in the spatial data, IPMN-estrogen pattern 5 is most highly weighted in LG IPMN as compared to HG and IPMN-PDAC (p < 2.2e-16, Kruskal-Wallis, p adjusted = 0 for LG vs. HG, p adjusted = 2.84e-49 for LG vs IPMN-PDAC, Dunn post hoc test with Bonferroni adjustment).

Similarly to IPMN-glycolysis pattern 3, IPMN-estrogen pattern 5 was highly weighted in a cluster containing both IPMN grades in the original cohort but may be more easily segregated by grade in higher resolution analysis. Given the smaller sample size of this cohort as compared to our original cohort, further analysis of these patterns in spatial data with single-cell or subcellular-level resolution should be performed to confirm their association with high or low grade IPMN, respectively.

We next examine whether there are significant differences in pattern weight by spatial location in reference to the lesion in epithelial spots (epilesional, or overlapping with neoplastic epithelium, juxtalesional, or surrounding the neoplastic epithelium, or perilesional, distal to the juxtalesional region). High-grade IPMN does not contain any annotated regional spots per the original authors, so we limit our consideration to low-grade IPMN and IPMN-PDAC. There are no observable differences in projected pattern weight by region for most patterns (**Supplementary Figure 3A-B).** There is a small yet statistically significant difference in IPMN-EMT pattern 1 weight between the epilesional and perilesional regions in IPMN-PDAC samples (p adjusted = 0.0216, ANOVA with Tukey post-hoc HSD) and across all samples regardless of tissue type (p adjusted = 0.0288, ANOVA with Tukey post-hoc HSD), with the pattern weight being higher further from the neoplastic epithelium **(Supplementary Figure 3B, Figure 3D)**. This may suggest that IPMN-EMT pattern 1, while identified and expressed in the epithelium, may also be exacerbated by increased cancer-associated fibroblast presence or other microenvironmental factors at a greater distance from the lesion itself.

To assess this hypothesis, we identify spatial regions, or “hotspots,” of inferred interaction between spots with enriched pattern expression and RCTD-annotated cell types using the SpaceMarkers tool^37^. We first illustrate this analysis in one representative sample (IPMN-PDAC #1039) in which we are able to spatially visualize areas of putative interaction between spots expressing a given transcriptional pattern and spots belonging to a particular cell type **(Figure 3E-G)**. In the IPMN-PDAC sample, areas of high projected IPMN-EMT pattern 1 weight overlap with both epithelial and fibroblast regions, as expected **(Figure 3E-F)**. We observe areas of interaction between IPMN-EMT pattern 1-expressing spots with fibroblasts in this sample, however the highest percentage in overlap of hotspots for this pattern is for various lymphoid and myeloid cell subtypes such as DC2.3 (annotated in the scRNA-seq reference atlas as CD1C^+^ dendritic cells), CD8^+^ memory T lymphocytes, and naïve CD4^+^ T cells **(Figure 3G, Supplementary Figure 4C).** CD1C^+^ dendritic cells are thought to induce the CD4^+^ helper T cell response for tumor antigen presentation^43^. CD4^+^ T cells are thought to play a role in EMT in the context of the pancreas, although the CD4^+^CD25^-^ T effector cell subtype most robustly demonstrated to induce EMT in pancreatic ductal epithelium^44^ is not present in this atlas. In contrast, IPMN-glycolysis pattern 3-expressing spots contain a much smaller area of interaction with fibroblasts, but relatively large areas of interaction with epithelial cells, overlapping with the entirety of the ductal epithelium, and with macrophage type 1 cells in the region of the duct itself **(Figure 3G)**. These macrophages were annotated in the single-cell reference atlas as expressing APOE and SPP1, as opposed to type 2 macrophages which expressed complement molecules^45^.

APOE+ macrophages have been demonstrated to affect immunotherapy outcomes in cancer through hypothesized interactions with CD8+ T effector exhausted cells^46^. It is possible that these macrophages interact with IPMN-glycolysis pattern 3-expressing epithelial cells to produce deleterious microenvironmental changes.

A broad look at all hotspots of interaction across all patterns and cell types across all samples in this dataset reveals distinct patterns of interaction between each pattern and annotated cell type by IPMN grade (LG, HG, or IPMN-PDAC) **(Supplementary Figure 4A-C).** For each IPMN grade, we compute the percentage of the total number of spots identified as belonging to a given pattern or cell type which are deemed as “hotspots”. In LG IPMN, we find that IPMN-EMT pattern 1 has the highest percentage interaction with fibroblasts, as expected. IPMN-glycolysis pattern 3-expressing spots exhibit the strongest interactions with NK and epithelial cells, while IPMN-estrogen pattern 5, IPMN-SR pattern 7, and pattern 9-expressing spots exhibit the strongest interactions with epithelial cells. Percentages of interaction with lymphocytes and myeloid cells are generally low across all patterns **(Supplementary Figure 4A).** In HG IPMN, the landscape of interactions changes dramatically, with a high percentage of interactions not only between IPMN-EMT pattern 1-expressing spots and fibroblasts and IPMN-estrogen pattern 4, IPMN-SR pattern 7, and pattern 9-expressing spots with the epithelium, as observed in LG IPMN, but also between each of the aforementioned patterns and T lymphocytes, dendritic cells, myeloid cells, and endothelial cells **(Supplementary Figure 4B)**. In IPMN-PDAC samples, the dominant interaction between IPMN-EMT pattern 1-expressing spots is with type 2 macrophages. While IPMN-estrogen pattern 5 and IPMN-SR pattern 7 both exhibit interactions with lymphocyte and myeloid subtypes, their dominant interaction is with the epithelium, similar to in LG IPMN **(Supplementary Figure 4C).** These results suggest that, in addition to possessing distinct gene expression programs and different levels of expression in each tissue type and/or region, each pattern has a distinct spatial distribution and cell type interaction profile which reveals microenvironmental heterogeneity of different IPMN grades not initially observed in our bulk RNA-seq analysis.

### Comparative analysis to PanIN reveals one IPMN-specific transcriptional phenotype and one pan-precancer phenotype

In the interest of determining which of our transcriptional patterns may be pan-precancer or pan-epithelial versus IPMN-specific, we project the CoGAPS pattern weights from the Semaan et al. RNAseq dataset onto a spatial transcriptomic dataset from a different type of pancreatic precancerous lesion, PanIN^14^. Spot annotations were obtained by the original authors via clustering and CODA, a machine learning approach to identify pancreatic cell types by morphology^47^. Epithelial spots were annotated as either normal, low grade PanIN, or high grade PanIN **(Supplementary Figure 5A)**. We apply the same pipeline above to determine the correspondence between our IPMN patterns in PanINs and use ProjectR to project feature weights of patterns of interest onto the PanIN query data (note that projected pattern weights less than zero represent the absence of the pattern in the query and thus are set to zero) **(Figure 4A, Supplementary Figure 5B)**. Interestingly, IPMN-glycolysis pattern 3, which was associated with KRAS c.436G>C and GNAS c.601C>A (from cluster IPMN-P3) in the original cohort, is not found in this data, neither in the dysplastic epithelium, normal epithelium, or nonepithelial regions. From this, we hypothesize that the transcriptional phenotype associated with IPMN-glycolysis pattern 3 is unique to IPMN and not generalizable to other types of pancreatic precancer such as PanIN. GNAS mutations in general are commonly observed in IPMNs but rare in PanINs, so this observation fits with the observed genetic profile of this phenotype in IPMNs^48^. We verify this hypothesis in a single-cell RNA-seq dataset from sporadic PanIN lesions in the adult pancreas^4^ where we also do not expect to find the unique mutational profile of the IPMN-P3 phenotype nor high weights of its characteristic IPMN-glycolysis pattern 3 within the epithelium. As hypothesized, IPMN-EMT pattern 1 (associated with cluster IPMN-PDAC) is found in all epithelial categories (adjacent normal, sporadic PanIN lesion, and normal epithelium), while IPMN-glycolysis pattern 3 not expressed in either normal, adjacent normal, or PanIN epithelium **(Figure 4B)**, indicating that it may be expressed only by epithelium with a particular IPMN-specific mutational profile.

**Figure 4.**
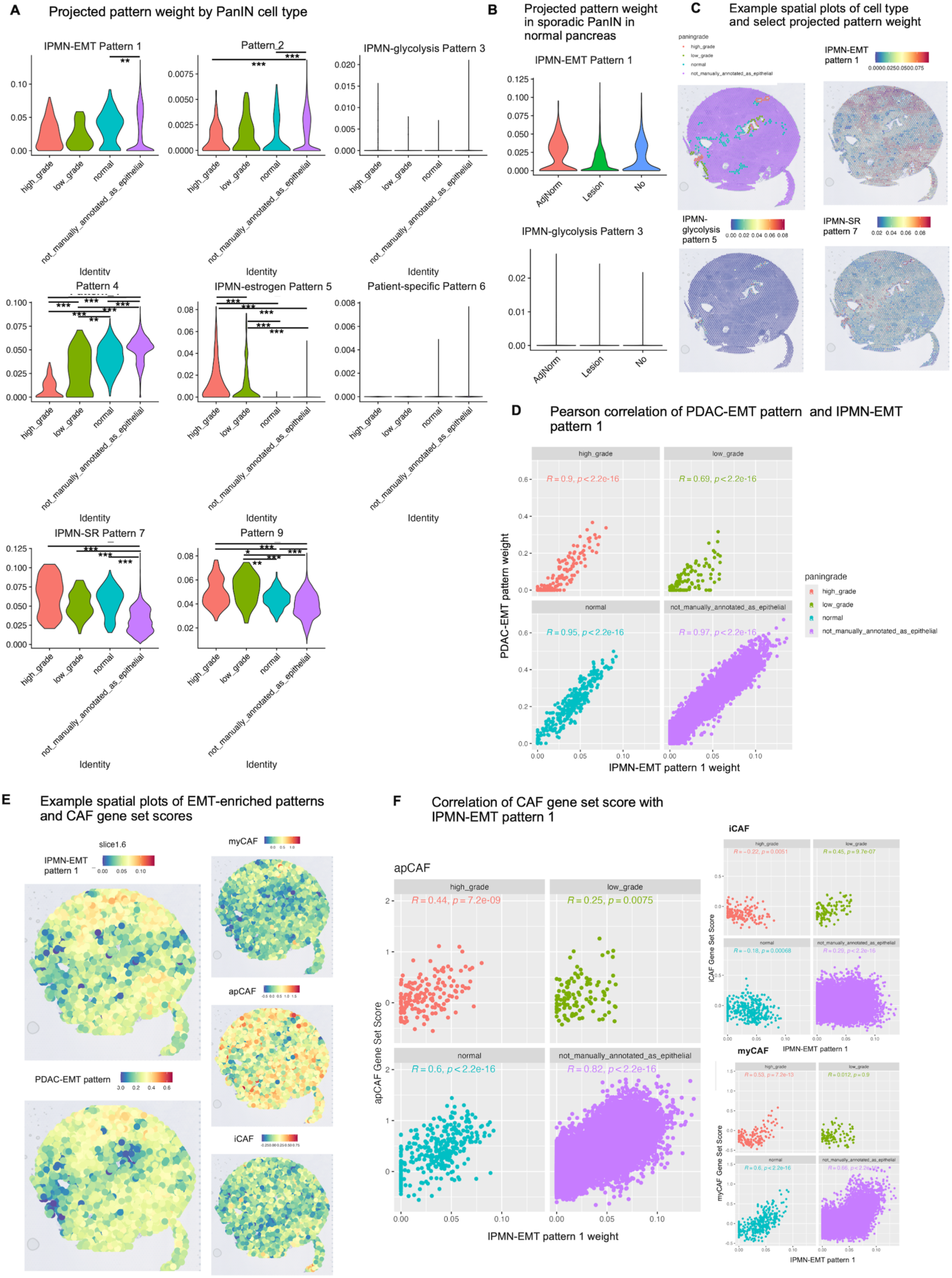
Comparative analysis to PanlN precancerous lesions

IPMN-estrogen pattern 5 and IPMN-SR pattern 7, which were both highly weighted in our IPMN-PDAC cluster, are enriched in dysplastic epithelium (IPMN-estrogen pattern 5) or epithelial cells as compared to nonepithelial cells (IPMN-SR pattern 7). IPMN-estrogen pattern 5 is highly weighted in high grade PanIN as compared with low grade PanIN, normal epithelium, and nonepithelial tissue (*p adjusted* < 0.0001 for all comparisons, Kruskal-Wallis with post-hoc Dunn and Bonferroni adjustment), and it is also enriched in low grade PanIN as compared to normal epithelium and nonepithelial tissue (*p adjusted* < 0.0001 for all comparisons, Kruskal-Wallis with post-hoc Dunn and Bonferroni adjustment). This pattern appears isolated to dysplastic epithelium and represents a transcriptional phenotype that becomes more dominant as PanIN lesions increase in grade of dysplasia. As hypothesized based on its weight across tissue types in the original cohort, IPMN-SR pattern 7 appears to be a pan-epithelial pattern with no difference in weight in normal versus dysplastic epithelium (p *adjusted* < 0.0001 for all comparisons, Kruskal-Wallis with post-hoc Dunn and Bonferroni adjustment). IPMN-EMT pattern 1, which was enriched in EMT pathways in IPMN, is found to be similarly weighted across dysplastic and normal epithelium in PanIN, with a significantly higher projected weight in nonepithelial tissue versus epithelial (*p adjusted* < 0.001, but no significant difference between dysplastic and nonepithelial tissues). We hypothesize that IPMN-EMT pattern 1 likely represents a combination of a stromal signal which contaminated the bulk RNA-seq data from nearby cancer-associated fibroblasts and epithelial-specific phenotype of expressed stroma-associated genes. We illustrate these pattern relationships spatially using sample 6, which is the only section containing both LG and HG IPMN **(Figure 4C)**. From this analysis, we confirm that IPMN-estrogen pattern 5 represents a transcriptional phenotype common to pancreatic precancer subtypes, namely the IPMN-PDAC cluster from the original cohort and high grade PanIN, while the *GNAS* mutation-associated IPMN-glycolysis pattern 3 is unique to IPMN.

In the original Bell et al. study^14^, patterns derived from a scRNA-seq atlas of advanced PDAC^12^ were projected onto the spatial PanIN data. The original authors found that two patterns emerged as correlated or anti-correlated with dysplastic progression: pattern 2, associated with KRAS signaling and proliferation and increasing in weight from normal epithelium to low and high grade PanIN, and pattern 7 (referred to here as the “PDAC-EMT pattern”), enriched in inflammatory and EMT signaling, correlated with cancer-associated fibroblast (CAF) density, and decreasing in weight as the grade of dysplasia increased^14^. We compare the PDAC-EMT pattern with IPMN-EMT pattern 1 and find that these two patterns are highly correlated in all tissue types: nonepithelial tissue (R = 0.97, p < 2.2e-16, Pearson correlation), normal epithelium (R = 0.95, p < 2.2e-16), low grade PanIN (R = 0.69, p < 2.2e-16), and high grade PanIN (R = 0.9, p < 2.2e-16) **(Figure 4D).** Upon visual inspection of the spatial location of these patterns, they also appear to co-localize **(Figure 4E**. We hypothesize that these EMT-enriched patterns represent a shared transcriptional phenotype in both IPMN and PanIN.

We also examine the relationship between IPMN-EMT pattern 1 and CAF subtype gene set scores defined in that original study. We obtain the gene set scores of three CAF subtypes as defined in the Bell et al. study: myofibroblastic CAFs (myCAFs), inflammatory CAFs (iCAFs), and antigen-presenting CAFs (apCAFs), where myCAFs and iCAFs arise early in PDAC development^14^. myCAF score is correlated with projected IPMN-EMT pattern 1 weight in all cell types except for low grade PanIN, apCAF score is correlated with IPMN-EMT pattern 1 weight in all cell types, and iCAF is anticorrelated with IPMN-EMT pattern 1 weight in high grade and normal epithelium but weakly correlated in low grade and nonepithelial tissue. Both EMT-enriched patterns appear to spatially co-localize with apCAF gene set score but not with myCAF or iCAF gene set score **(Figure 4F).** We hypothesize that IPMN-EMT pattern 1, while expressed in epithelial cells, is also strongly associated with CAF presence, specifically the apCAF subtype. These results demonstrate that some patterns, such as IPMN-EMT pattern 1 which was associated with the IPMN-PDAC cluster, may be related to microenvironmental differences while other patterns, such as IPMN-glycolysis pattern 3 which corresponds to the IPMN-P3 cluster in the IPMN atlas, may be unique to IPMN-specific mutational signatures.

To extend our analysis beyond CAFs and compare the microenvironmental effects in IPMN to PanIN lesions, we identify hotspots of interaction between author-defined cell types and projected IPMN-derived patterns as with the previous spatial dataset. The original authors quantify percentages of each cell type per spot using the H&E-based cell type reconstruction method, CODA^47^. Cell types include acini, collagen, fat, islets, normal epithelium, PanIN epithelium, and smooth muscle. Of note, spots containing no cells are annotated separately, and the “collagen” spots are collagen-rich on H&E and express a panCAF signature^14^. Patterns with the highest relative interactions with collagen-rich spots, presumably containing CAFs, include IPMN-EMT pattern 1 as expected (in samples LG1, LG2, HG1, and HG4) and pattern 4 (in sample LG3). IPMN-estrogen pattern 5 displays the highest relative interaction with PanIN epithelium as opposed to normal epithelium across all samples but one (sample LG1, LG3, HG1, HG2, HG3, and HG4). We observe some individual heterogeneity in interactions between pattern-expressing cells and broad cell types, however the patterns with the relatively greatest interactions for a given cell type are well preserved across samples and grades of dysplasia **(Supplementary Figure 5D)**. These results suggest that IPMN-EMT pattern 1 is not only correlated with CAF gene expression but may represent an epithelial-fibroblast interaction, and that IPMN-estrogen pattern 5 may be involved in dysplastic mechanisms within the epithelium and may represent a marker of progression to PDAC.

### Pancreatic precancer datasets and IPMN-derived patterns are made publicly available for user-friendly analysis

The analysis in this study enables cross-comparison of transcriptional phenotypes throughout pancreatic precancer lesions and into advanced pancreatic cancer. While powerful, the analyses performed required us to collate each of these datasets and manually compare them through bioinformatics scripts. The Gene Expression Analysis Resource (gEAR) portal^19, 49^ provides a user-friendly, web-based environment for bioinformatics analysis of multi-omic data. Of relevance to our study, it includes the ability to visualize gene expression and to perform transfer learning analysis using the projectR tool of a transcriptional pattern of interest onto a query dataset. To enable users to perform cross-comparison of our IPMN-derived transcriptional programs and to interrogate the datasets analysed in this study, we have uploaded both the patterns and the spatial IPMN and PanIN datasets to Cancer gEAR. Our precancer datasets and gene signatures are fully available from https://cancer.umgear.org/landing/pdac/ and demonstrated through a schematic in **Figure 5**.

**Figure 5.**
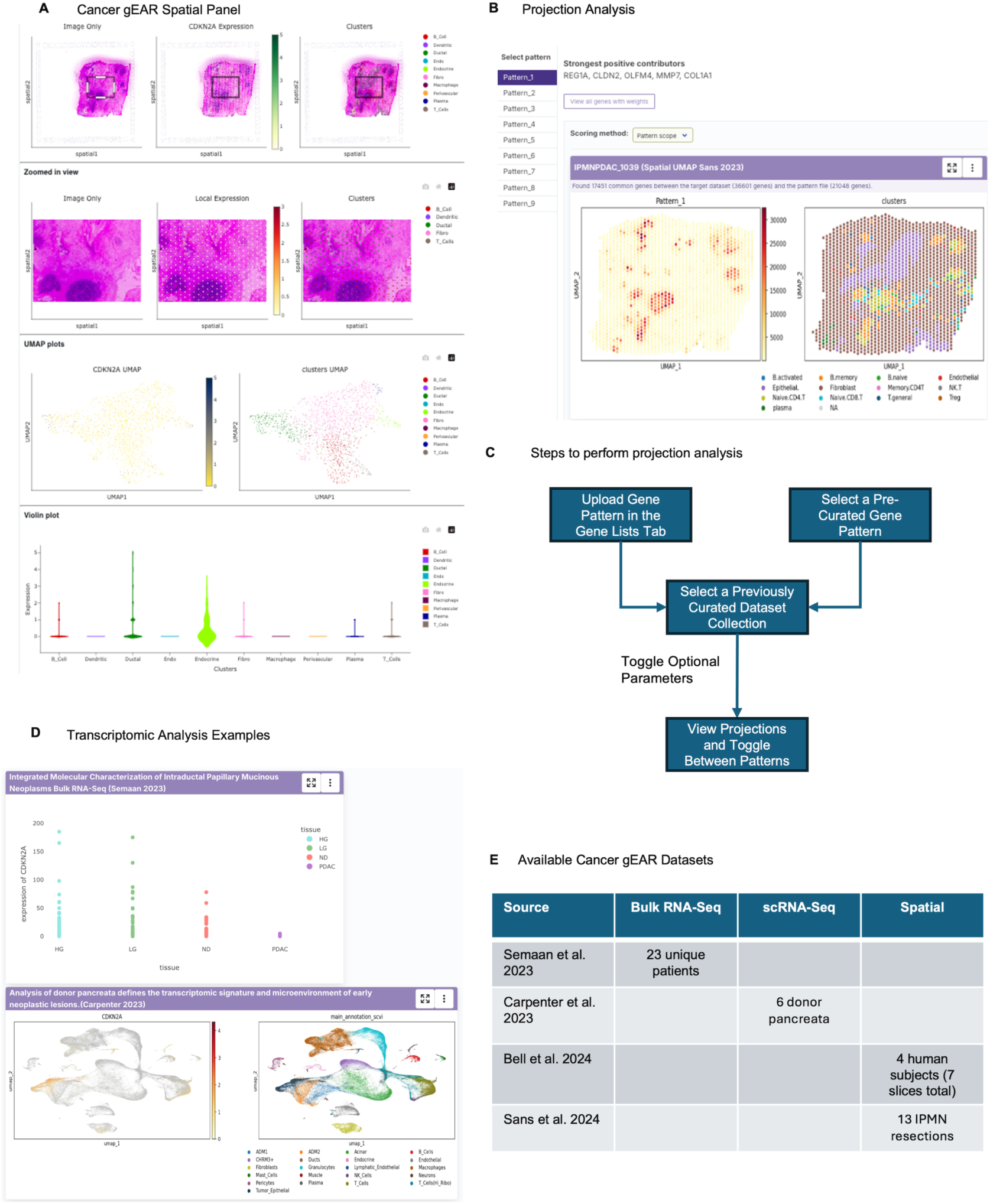
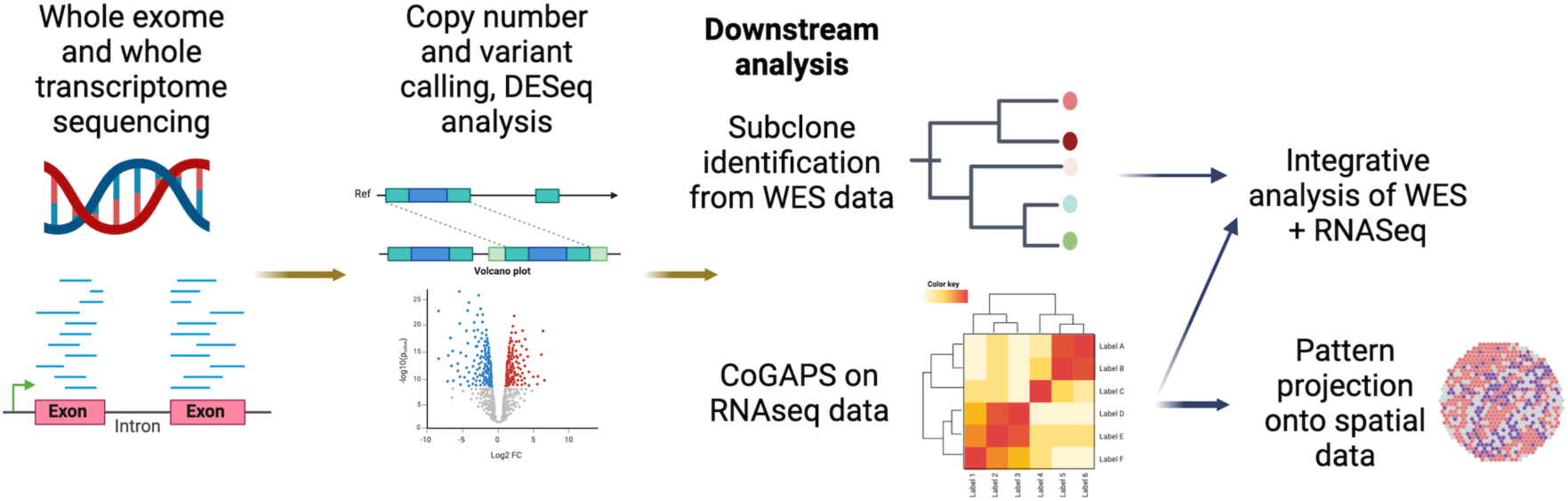
Public dissemination and user-friendly analysis and visualization enabled by the cancer gEAR platform

## Discussion

The goal of this study was to perform subtyping of IPMNs to better understand the mechanistic transformation of IPMNs into PDAC and to identify which subtypes are likely to progress versus remain indolent. Our findings highlight the challenge of relying on somatic mutation information alone for IPMN subtyping: as expected, we observed significant population-wide genomic heterogeneity with limited overlap of individual mutations across patients, which limited the ability to define generalizable subclones on a cohort-wide rather than a per-patient basis. Similarly, applying the established PDAC transcriptional subtypes resulted in the classification of all samples as “classical” according to clinically correlated subtypes from Moffitt et al.^8^. By leveraging non-negative matrix factorization for *de novo* pattern detection, we identified transcriptional programs that transcend patient-specific variation and are enriched in different tissue types, revealing cohort-wide transcriptional heterogeneity within IPMNs. Integrating these transcriptional phenotypes with genomic data uncovered distinct multi-omic IPMN phenotypes, some of which may be more strongly driven by mutational profile, while others are more strongly shaped by the microenvironment. Interestingly, none of the transcriptional patterns or multi-omic phenotypes in this study segregated the samples by dysplastic grade.

We also examine how the transcriptional patterns associated with each multi-omic subtype appear in a spatial context in both IPMN and PanIN. The IPMN-PDAC multi-omic phenotype, which includes low-grade, high-grade, and PDAC samples from the original cohort, is comprised of three transcriptional programs, each of which appear in PanIN and one of which (IPMN-estrogen pattern 5) increases in weight in PanIN epithelium as it progresses from low-grade to high-grade dysplasia. We hypothesize that the estrogen pathway signature relates to non-canonical signaling and cellular proliferation and growth involving the c-Myc pathway, which has been hypothesized to play a role in PDAC progression^50^. Another pattern associated with this phenotype, IPMN-EMT pattern 1, appears to be associated with fibroblast presence and epithelial-fibroblast interactions in IPMN and PanIN and is positively correlated with a PDAC-derived EMT transcriptional program in PanIN and is independent of the Moffitt et al. basal subtype commonly associated with EMT^8^.

This IPMN-PDAC cluster is an example of a phenotype strongly influenced by microenvironmental features and present across different types of pancreatic precancer as well as PDAC. By contrast, phenotype IPMN-P3 is named as such because it is strongly associated with expression of a single transcriptional pattern, IPMN-glycolysis pattern 3, which does not appear in PanIN upon cross-comparison. This phenotype also presents a unique somatic mutation profile to the IPMN-PDAC phenotype, containing a different set of *KRAS* and *GNAS* mutations, however due to the limited sample size of the WES data in the original cohort this somatic mutation profile cannot be generalized without further comparative analysis. A further limitation of this study arises from the lack of mutational data for our PanIN cohort, hindering our ability to associate the transcriptional phenotypes with driver mutations or mutational subtypes in this type of precursor lesion.

Transfer learning analysis of the phenotype-associated transcriptional patterns in spatial IPMN and PanIN data also presents the opportunity to estimate cell-cell interactions between pattern-expressing cells and various cell types of interest. Our hotspot analysis in IPMN spatial data reveals that pattern-expressing cells interact with different sets of stromal or immune cells depending on dysplastic grade. Quantification of the relative area of interaction between a given transcriptional pattern and cell type reveals that the patterns associated with a given multi-omic phenotype do not necessarily interact with the same cell types, revealing the multi-layered contribution of microenvironmental influence to transcriptional programming in these samples. In a hotspot analysis of PanIN data, the same patterns are found to have predominant interactions with the same broad cell types across the majority of samples, suggesting that some pan-precancerous patterns may interact with the same cell types regardless of dysplasia grade in PanINs. Further analysis of these hotspots of interaction in a spatial dataset with single-cell resolution is needed to confirm these findings.

To encourage comparative analysis of the transcriptional patterns identified in this study, we have made these patterns available to a user-friendly, web-based platform with a built-in tool for transfer learning analysis in any available query dataset on the platform. We have developed a section of this platform, termed Cancer-gEAR, dedicated to pancreatic precancer and cancer. The spatial transcriptomic precancer datasets discussed in this paper, which are already available on GEO, have now been uploaded to gEAR to enable visualization and analysis. This resource will enable other researchers with varying degrees of familiarity in R to perform comparative analyses of our gene signatures and the datasets used in this paper. Uploading these data are the first step towards building a pancreatic cancer and precancer-specific version of Cancer gEAR, similar to gEAR^19^ for the hearing community and NeMO Analytics^49^ the gEAR offshoot for neurobiological datasets. We hypothesize that further exploration will reveal associations and features of these patterns which will be productive for understanding the dysplastic progression and pathogenesis of pancreatic precancer and PDAC.

## Acknowledgements

We recognize funding from NIH/NCI (U54CA274371 to EJF, LDW, AM, MPM, and RK; U01CA271273 to EJF, LDW, RK; T32CA154274 to KN; P30CA134274 to AM, MG; U01CA200468 to AM; U24CA274274 to AM), the Lustgarten Foundation (to LDW), Sol Goldman Pancreatic Cancer Research Center at JHU (to LTK and LDW), and the Rackham Predoctoral Fellowship from the University of Michigan (to AE). Software development for the gEAR platform is supported by NIH/NIDCD (DC019370 to AM) and NIH/NIMH (R24MH114815 to AM, OW, MG).

We would like to thank Atul Deshpande for his input on SpaceMarkers analysis.

---

**Supplementary Figure 1.**
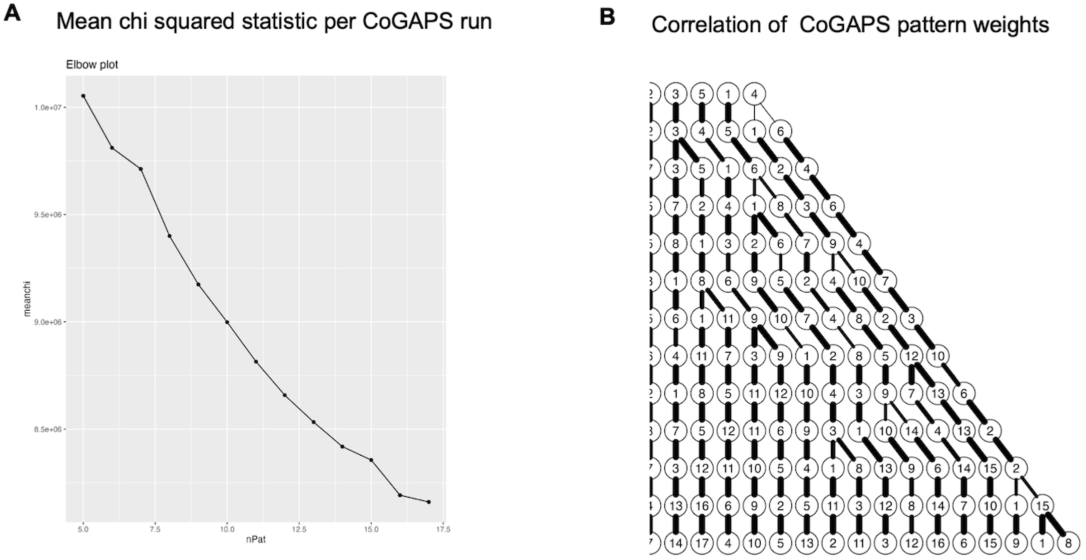
CoGAPS optimization

**Supplementary Figure 2.**
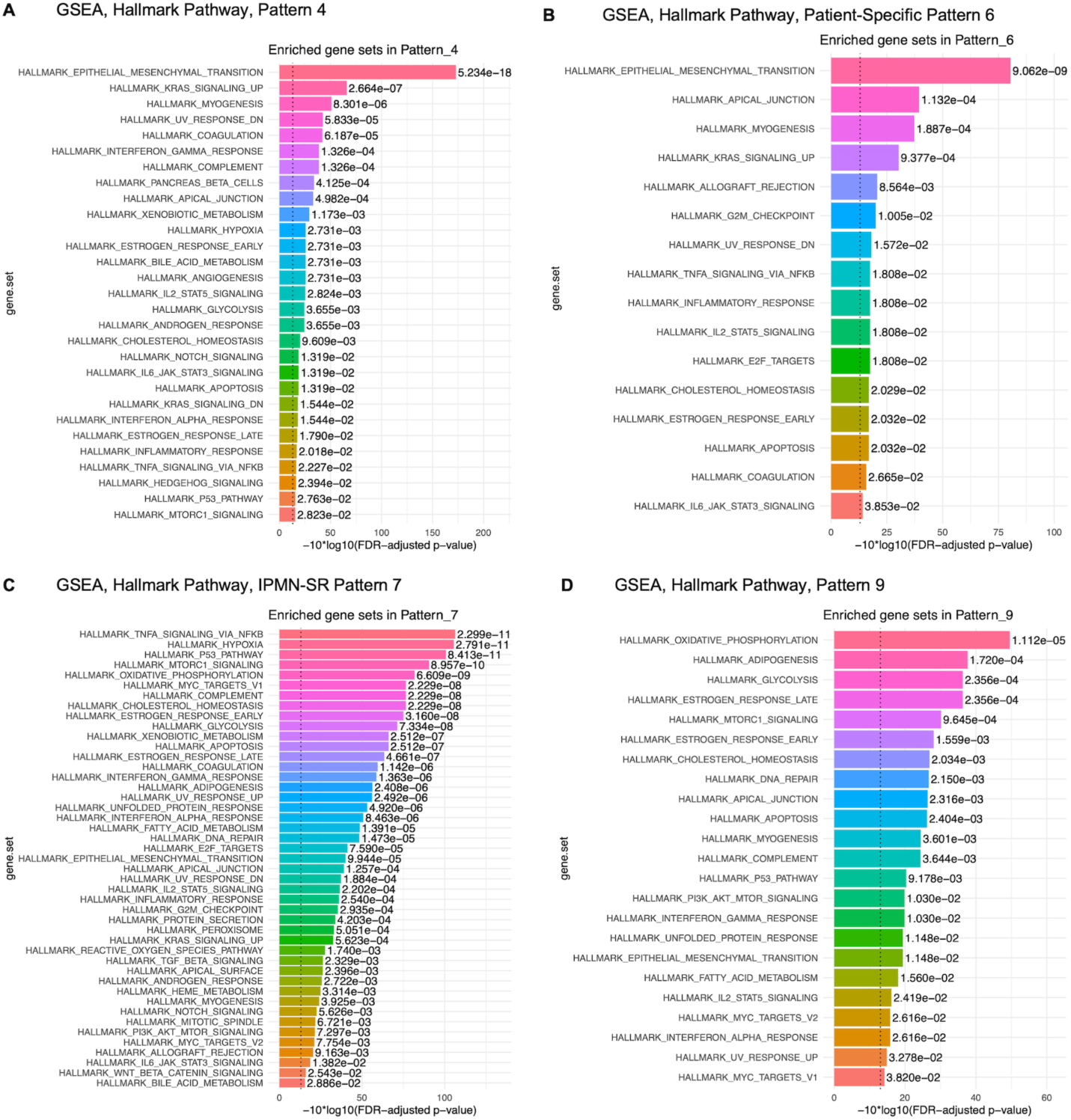
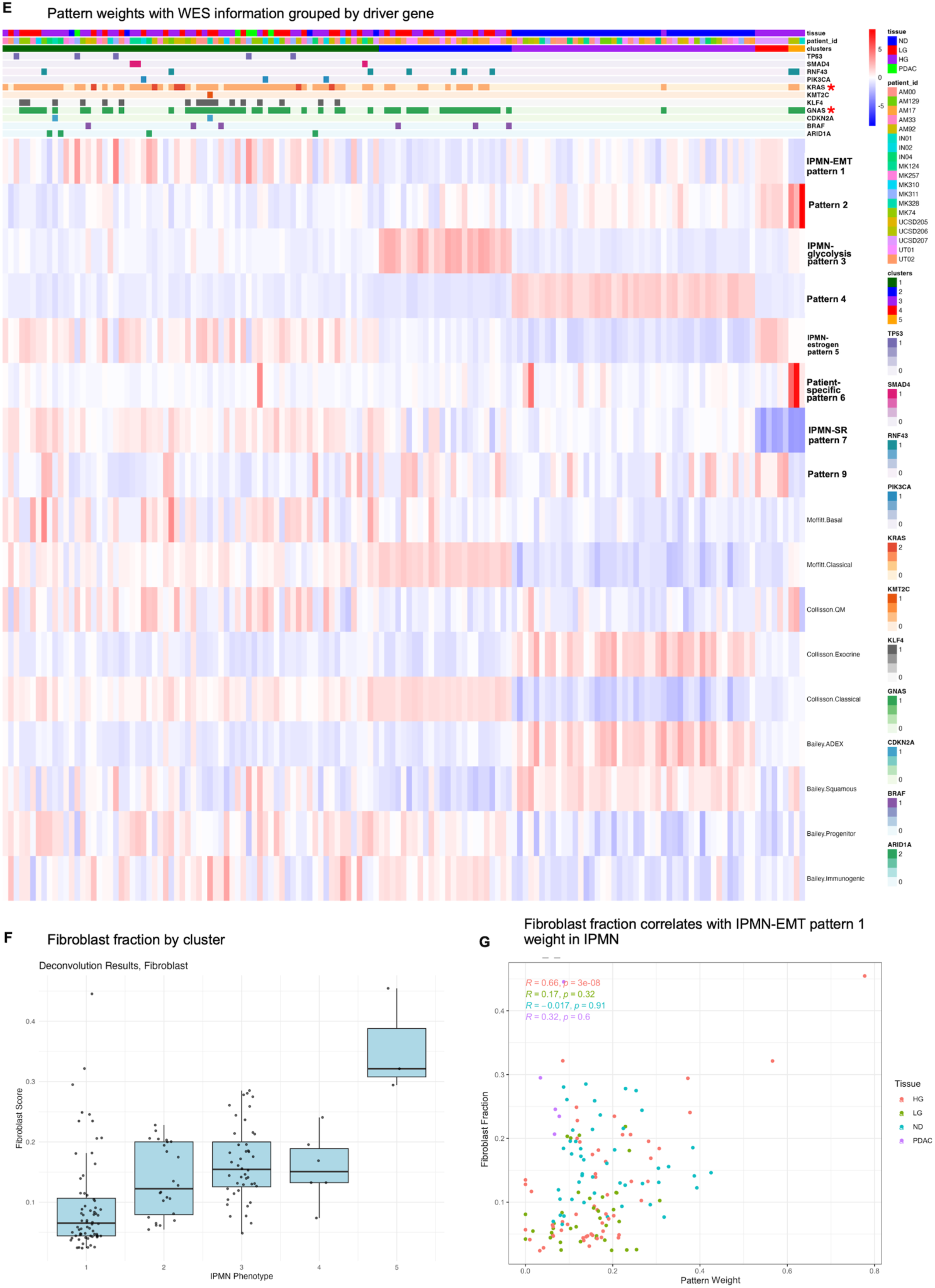
GSEA of remaining CoGAPS patterns

**Supplementary Figure 3.**
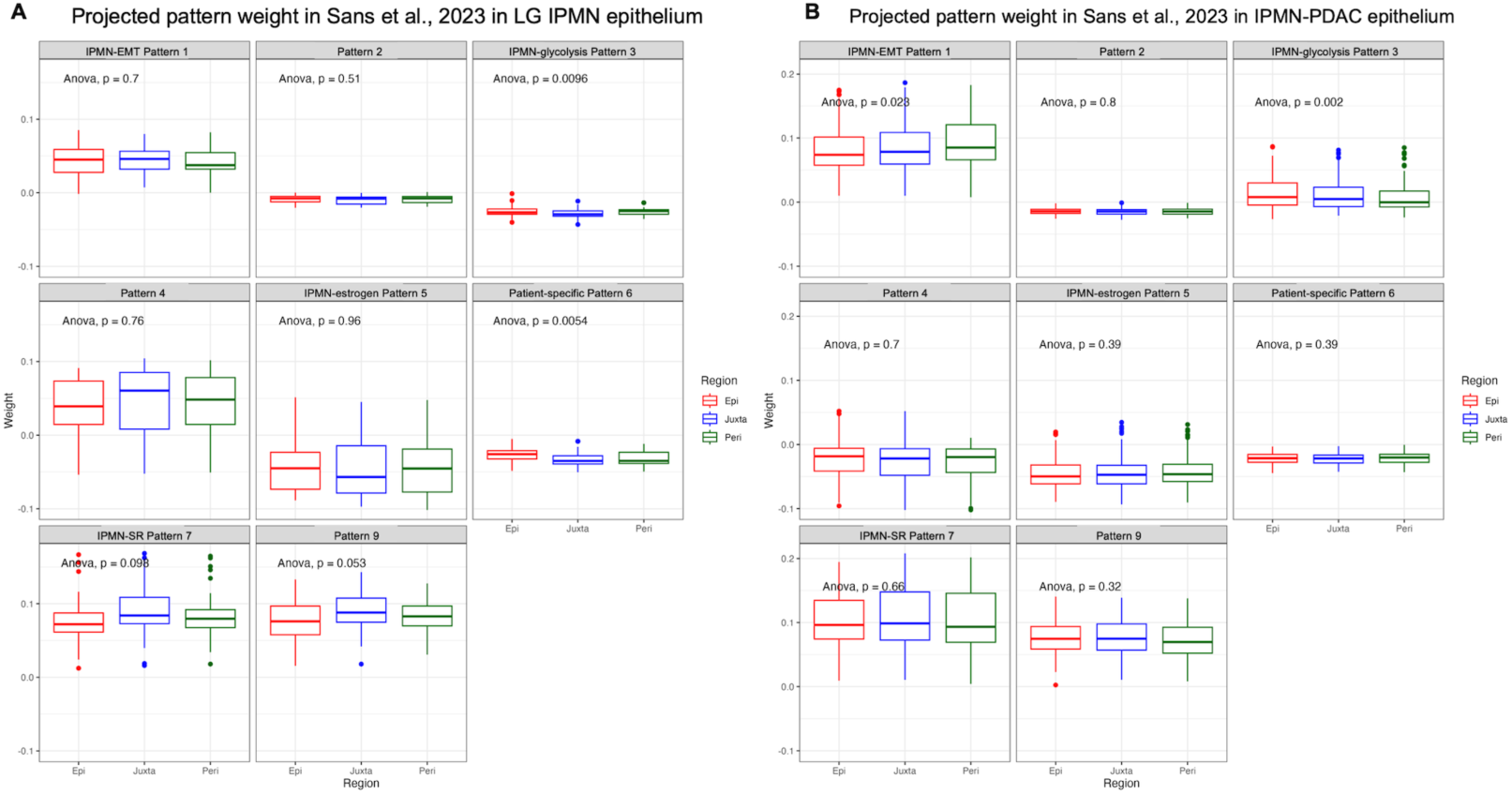
IPMN cross-comparison

**Supplementary Figure 4.**
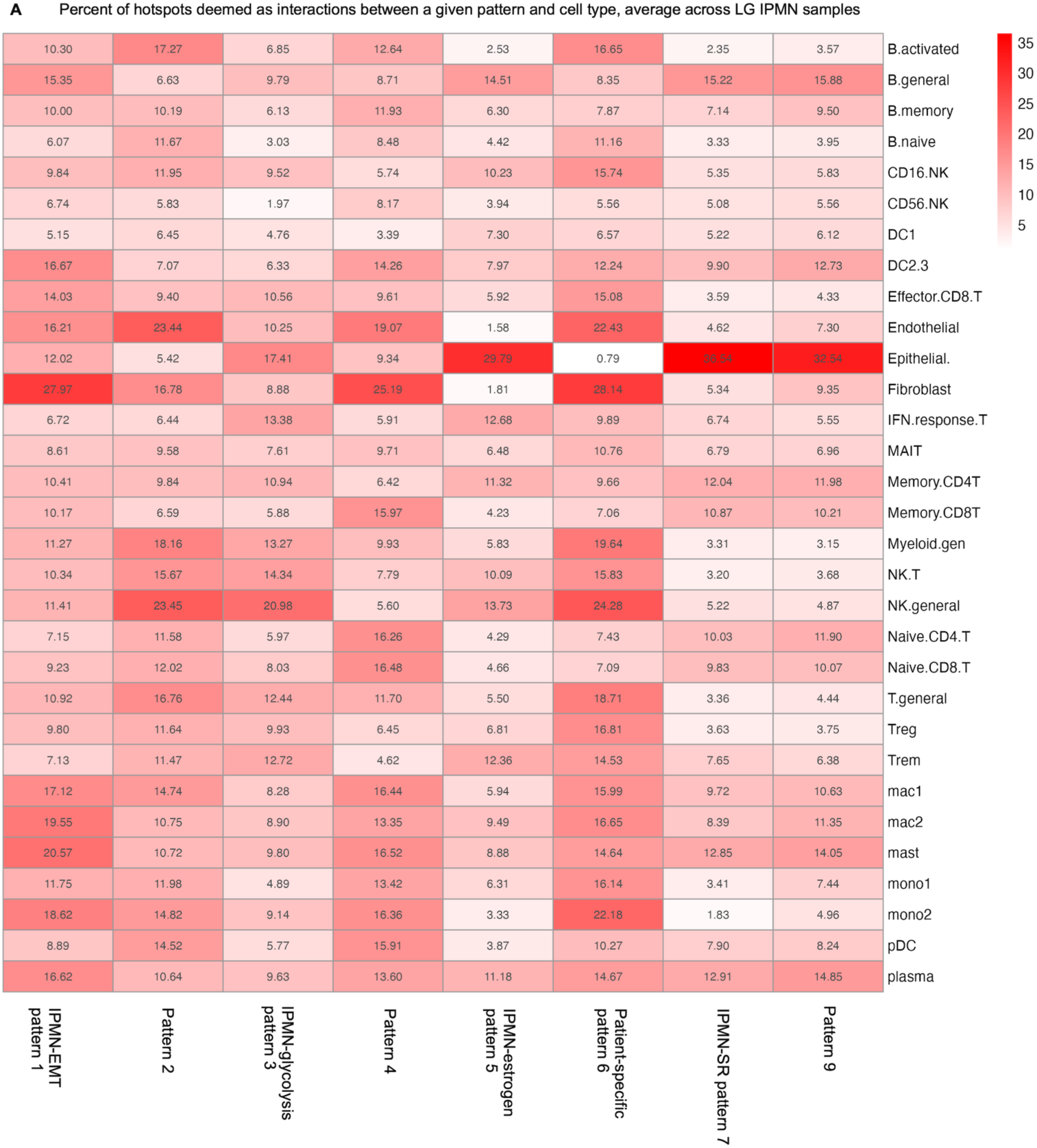
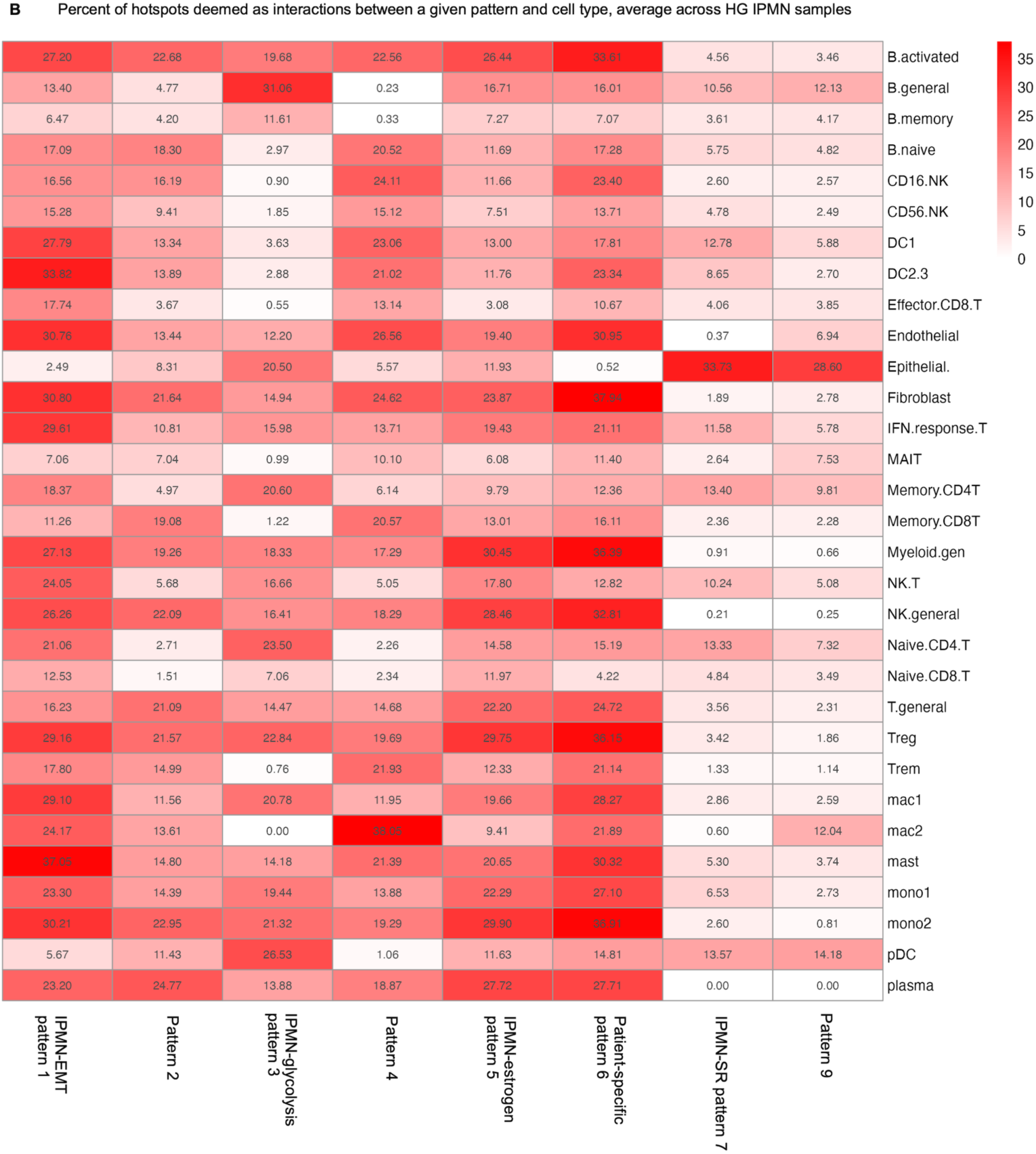
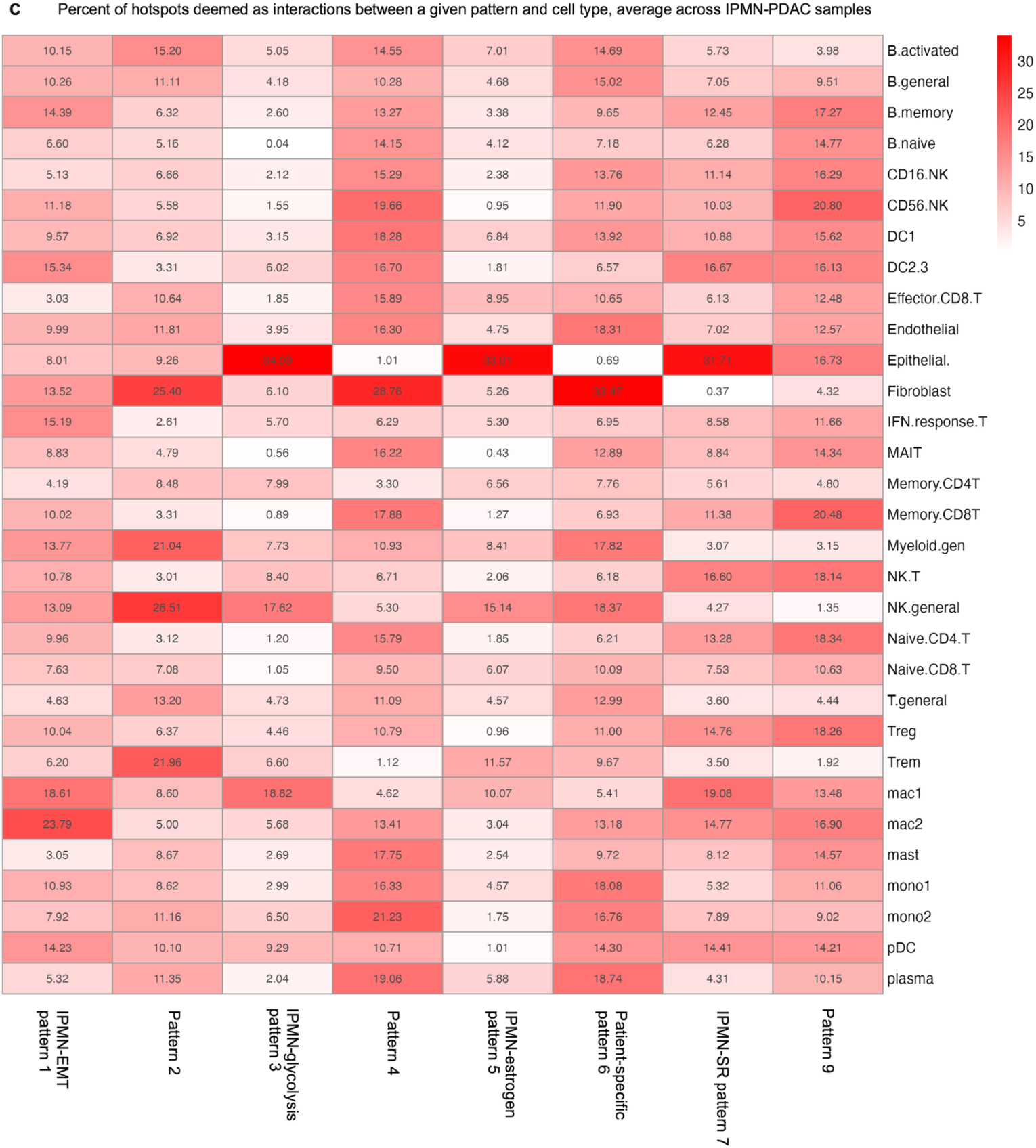
IPMN hotspot analysis

**Supplementary Figure 5.**
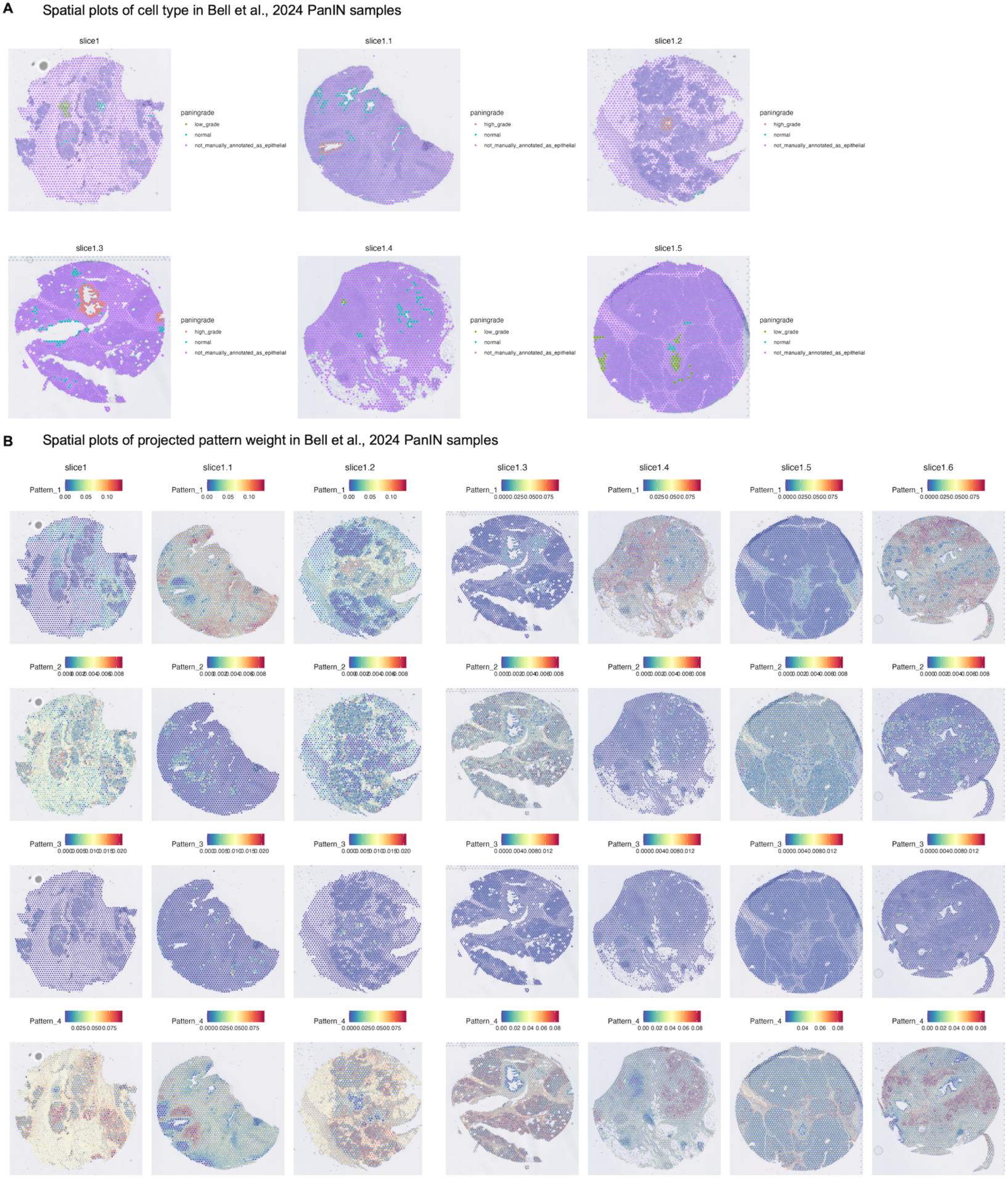
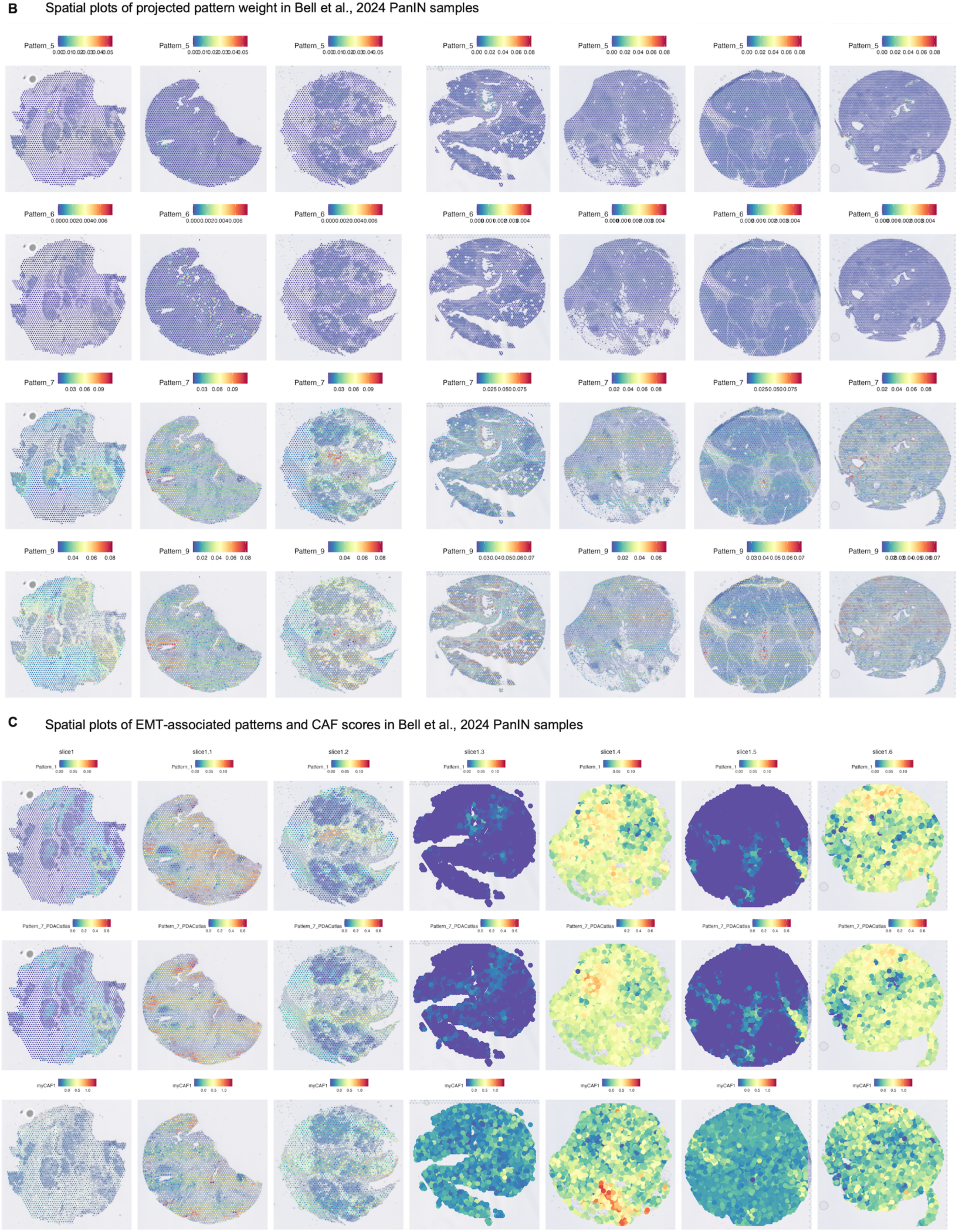
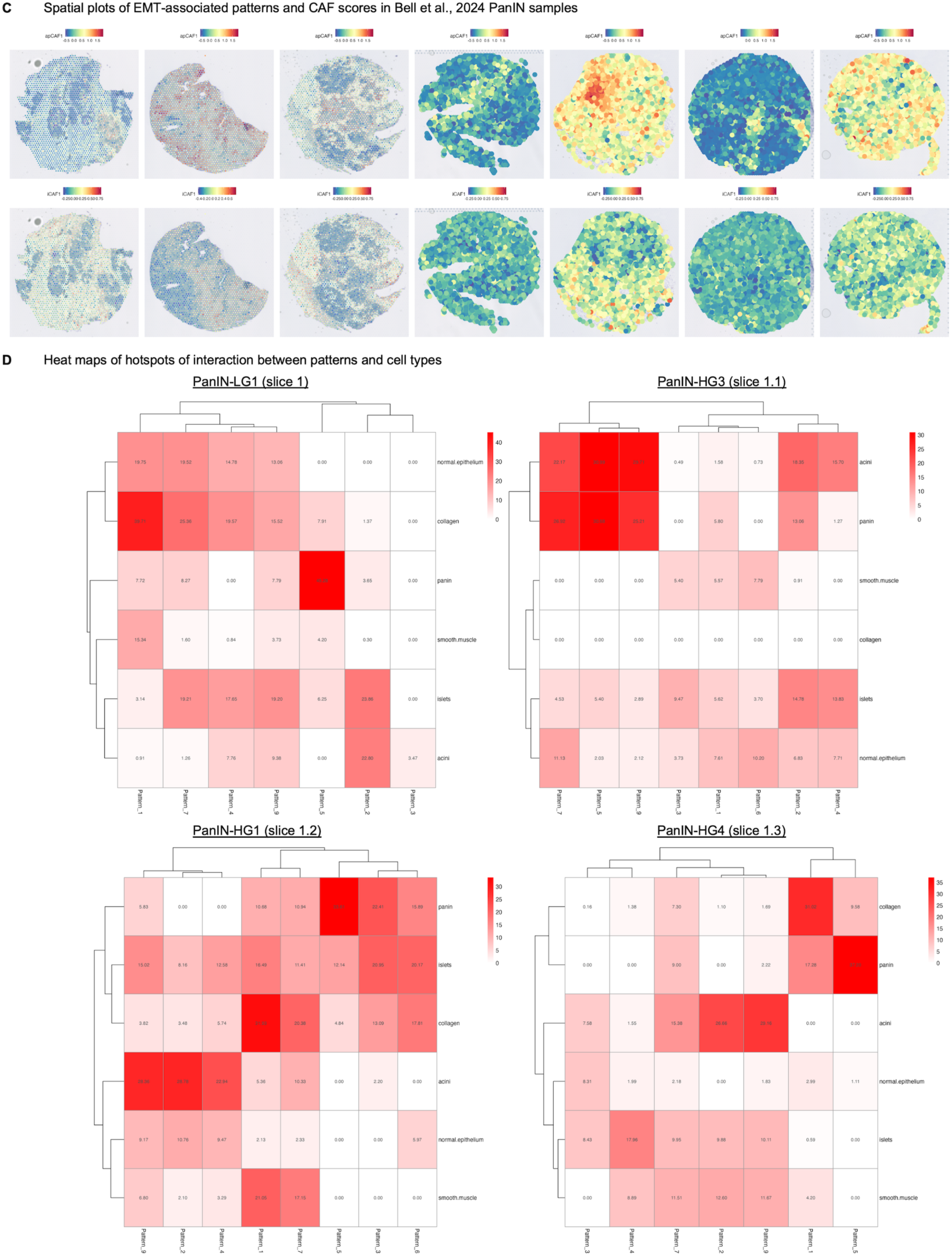
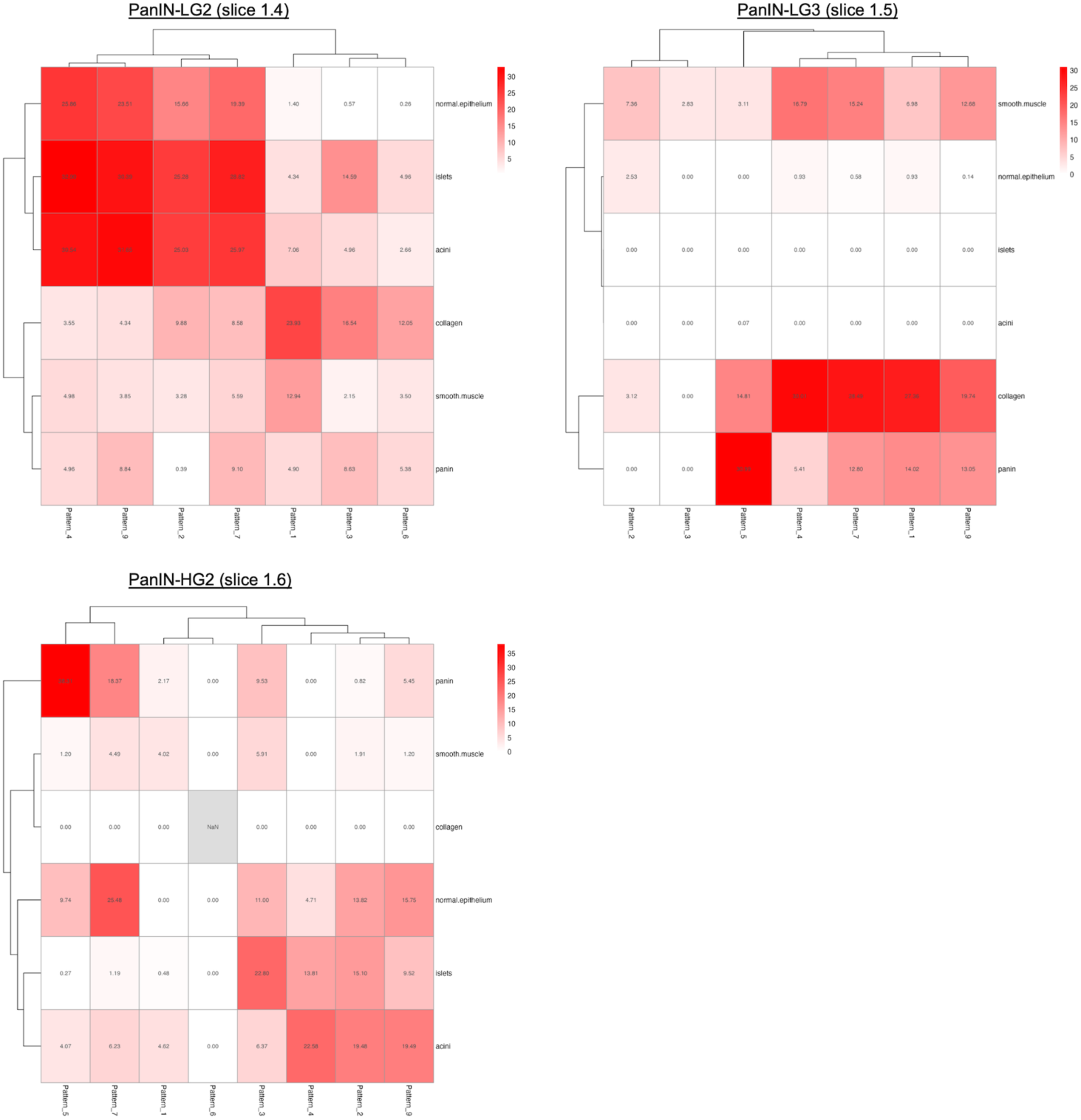
PanlN comparative analysis

